# Synaptic and potential fluctuations drive representational drift with differential effects on discrimination learning

**DOI:** 10.1101/2025.09.30.679426

**Authors:** Kento Nakamura, Keita Endo, Hokto Kazama

## Abstract

Large-scale neuronal measurements have revealed continual drift of neural representations without explicit learning, raising fundamental questions about how the brain maintains reliable perception. While representational drift has been studied both experimentally and theoretically, the underlying mechanisms and computational consequences remain unclear. Here, we systematically compare two drift mechanisms—fluctuations in synaptic weights versus membrane potential—using an analytically tractable neural network model with aligned drift magnitude and timescale. We found that both mechanisms preserve representational structure across time regardless of the drift magnitude. However, only synaptic fluctuations maintain discriminability between representations under large drift magnitude. We use a selectivity space to explain how these similar and differential effects on representational drift emerge from the dynamics of individual neuronal tuning. We further examine functional consequences of drift using reversal learning tasks where stimulus-reward associations switch. Both mechanisms enhance single stimulus learning by gradually removing the old associations, but only synaptic fluctuations substantially improved learning of multiple stimuli by preserving the representational structure during drift. Our results reveal distinct computations performed by specific fluctuation mechanisms and explain how representational drift can support both stable and adaptive behavior.

## Introduction

Neural representation of an external environment is expected to remain stable over time to support stable identification of stimuli. However, multiple studies have shown that neural representations can exhibit substantial changes over time. An early study reported that the place field of hippocampal pyramidal neurons can change even without explicit learning [1]. Recent advances in large-scale recording have revealed such instability also at the neuronal population level [2, 3, 4, 5, 6]. This dynamic reconfiguration, occurring without apparent learning demands, is actively discussed under the term “representational drift” [5, 6, 7, 8, 9]. The phenomenon has been observed in cognitive [10, 11, 12, 13], motor [14, 15], and even primary sensory regions [16, 17, 18, 19], raising fundamental questions about how and why drift emerges and how the brain maintains its reliable function despite drift.

Theoretical models have attempted to reproduce drift by incorporating stochastic fluctuations in synaptic strength [20, 21, 22, 23, 24, 25] or neuronal excitability [20, 21, 23, 26, 25] (see [9, 27, 28, 29, 30, 31] for other approaches), reflecting experimental evidence of changes in spine properties [32, 33, 34, 35] and excitability [36, 37, 38]. These studies have shown that even under substantial drift, networks can preserve informative structures such as assemblies [20, 23] or representational similarity [24, 30], and that unsupervised Hebbian mechanisms can alter readouts to stabilize outputs [22, 26]. However, most prior work has focused on a single source of fluctuations, making it difficult to determine how distinct mechanisms uniquely shape drift. Even when multiple sources were considered, their individual contributions were not systematically compared, leaving open the question of whether synaptic and excitability fluctuations induce qualitatively different forms of representational drift and therefore have differential downstream consequences [20, 21, 23, 25].

Here, we use an analytically tractable model of representational drift to address this question. We construct two models with fluctuations—one driven by synaptic fluctuations, the other driven by potential fluctuations— and calibrate them to exhibit identical magnitudes and timescales of drift. This alignment ensures that any residual differences in the network behavior should arise from the qualitative difference between drift mechanisms. In this framework, we find that both mechanisms preserve the representational structure across time in line with several experimental studies [13, 16, 39] (although see [17]). However, only synaptic fluctuations maintain discriminability between representations under large drift. Our analysis in a selectivity space provides a mechanism underlying the stability and instability of representations caused by the two fluctuations. Furthermore, we examine the consequences of distinct drift mechanisms on associative learning. Using a reversal learning task where stimulus-valence pairing occasionally changes, we find that both mechanisms similarly enhance reversal learning of a single stimulus-valence pair by gradually weakening old memory. However, in case of learning multiple stimulus-valence pairs demanding sensory discrimination, only synaptic fluctuation improves the performance.

## Result

### Model of representational drift

We used a simple two-layer feedforward network in which fixed input patterns were projected through random connections to a large hidden layer (Fig. 1A). This generates neurons with diverse tuning, similar to those observed in olfactory, cerebellar, and hippocampal circuits [40]. Hidden neurons are activated when the input current exceeds a threshold. Representational drift in the hidden layer was implemented either by adding stochastic fluctuations to synaptic connections or directly to the input current flowing into hidden neurons. At the level of abstraction of our dimensionless model, fluctuations added directly to the input current can be interpreted as fluctuations in the resting membrane potential. We thus term the former and the latter types of fluctuations as synaptic and potential fluctuations. We modeled fluctuations as a zero-mean Ornstein–Uhlenbeck (OU) process as in previous drift models [22, 26], which provides stationary variability with a characteristic magnitude (*σ*) and rate (*γ*) (or equivalently, timescale (*τ* := *γ*^−1^)) (see Methods). We note that such continual and activity-independent fluctuations are observed in the properties of synaptic spines [32, 33, 34, 35] and in neuronal excitability [37, 38]. To align the magnitude of drift under both mechanisms, synaptic fluctuations were scaled by the number of inputs to each neuron. To examine the changes in representation under constant drift with the same property, the fluctuating components of synaptic weights and input currents were sampled from the steady-state distribution of the OU process at the beginning of each simulation (time *t* = 0) and subsequently evolved according to the OU process dynamics. The activation threshold was set to make a certain fraction of neurons active at time *t* = 0 (see Methods).

**Figure 1:**
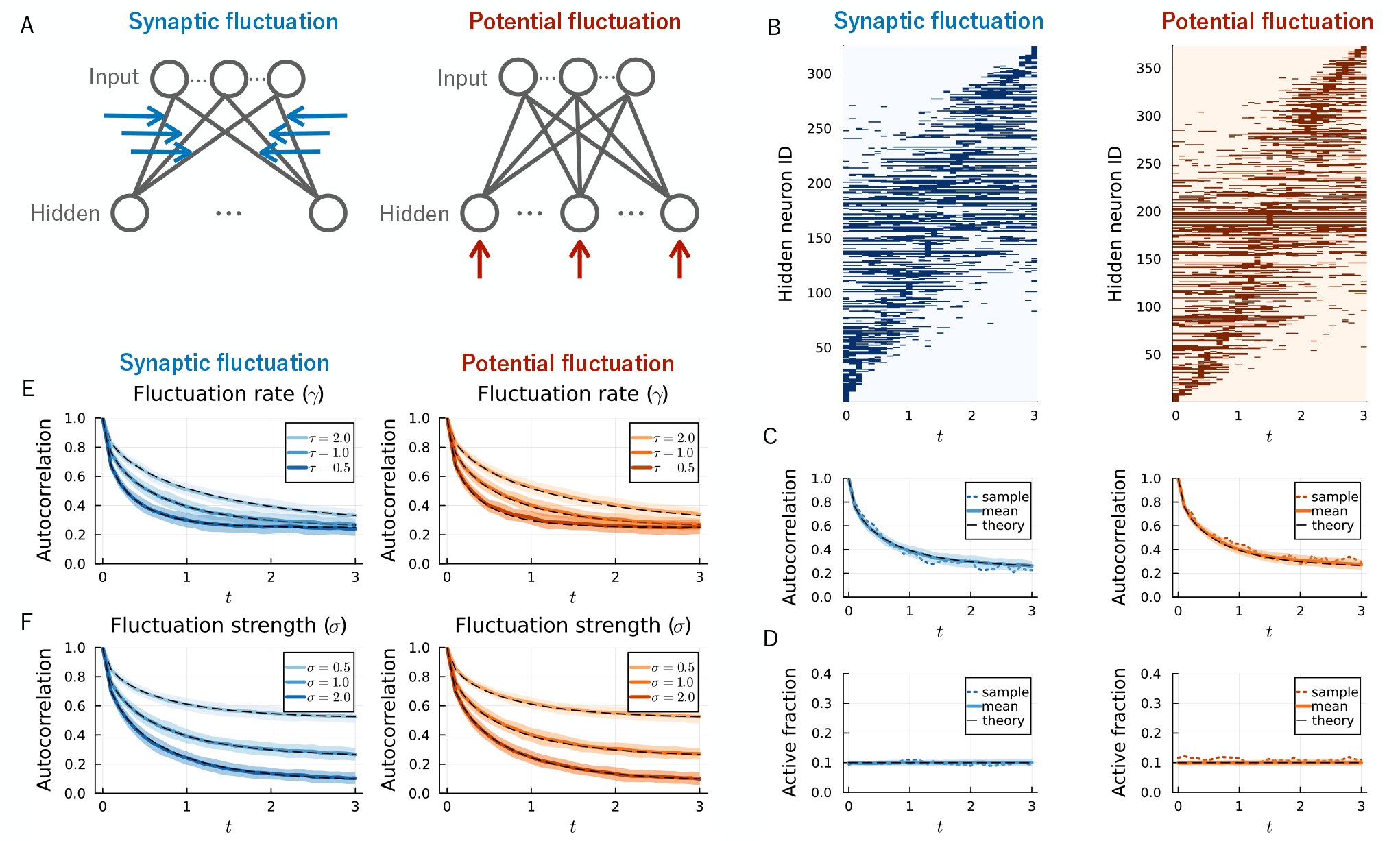
Drift model and autocorrelation of representations. (A) Network model of representational drift with two distinct mechanisms: synaptic fluctuation and potential fluctuation. The model consists of an input layer, a hidden layer, and random feedforward connections between layers. Synaptic fluctuations are added independently to each synaptic connection (blue arrows), whereas potential fluctuations are added independently to each hidden neuron (red arrows). (B) Sample trajectory of neuronal activity shows gradual representational drift for both synaptic and potential fluctuations. Hidden neurons that were active at least once in response to repeated stimuli are shown. (C,D) Autocorrelation (C) and fraction of active neurons (D) over time calculated using all neurons including non-responsive ones. Both mechanisms show gradual decorrelation over time while maintaining constant population sparseness. See Fig. S2 for results excluding non-responsive neurons. (E,F) Dependence of autocorrelation on fluctuation rate *γ* (E) and strength *σ* (F) demonstrates that the drift rate and magnitude are quantitatively aligned between the two mechanisms. In panels (C-F), solid curves and shaded regions indicate mean and standard deviation over 100 simulations, and dashed curves indicate theoretical predictions assuming infinitely large networks (Eqs. (1)-(4)). In panels (C,D), dotted, colored curves indicate a specific realization corresponding to the sample trajectory shown at the top. Parameters of the network were *γ* = 1.0, *σ* = 1.0, unless otherwise indicated in each panel.

### Representational drift under two different mechanisms

We examined how representations of a single stimulus drifted under synaptic and potential fluctuations. For both mechanisms, we observed that hidden neurons occasionally gained or lost activity over time, resulting in gradual drift of the population representation away from its initial state (Fig. 1B). This drift is reflected in a smooth decrease of autocorrelation over time (Fig. 1C). Despite ongoing drift, the overall fraction of active hidden neurons remained stable without adjustment of the activation threshold (Fig. 1D), consistent with analytical predictions (see Methods).

Representational drift under both synaptic and potential fluctuations showed similar dependence on the fluctuation parameters. The fluctuation rate *γ* controlled the speed of decorrelation (Fig. 1E), while the strength *σ* set the asymptotic level of autocorrelation (Fig. 1F). Although autocorrelation converged to a positive value at moderate *σ*, representations continued to drift while maintaining the same correlation level (Fig. S1B). Because the drift magnitude and timescale were matched between the two mechanisms, any qualitative differences observed later can be attributed to the nature of fluctuations rather than trivial differences in the rate or magnitude.

### The structure of representation is preserved across time during drift under both mechanisms

We examined the impact of synaptic and potential fluctuations on the relationship between representations of multiple stimuli. We considered a set of stimuli organized in a ring structure as a simplest example (Fig. 2A) (see also Ref. [24]). We constructed the input layer representations such that individual neurons tuned to specific phases tile the entire phase of the ring as a population (Fig. 2B; see Methods for detail).

**Figure 2:**
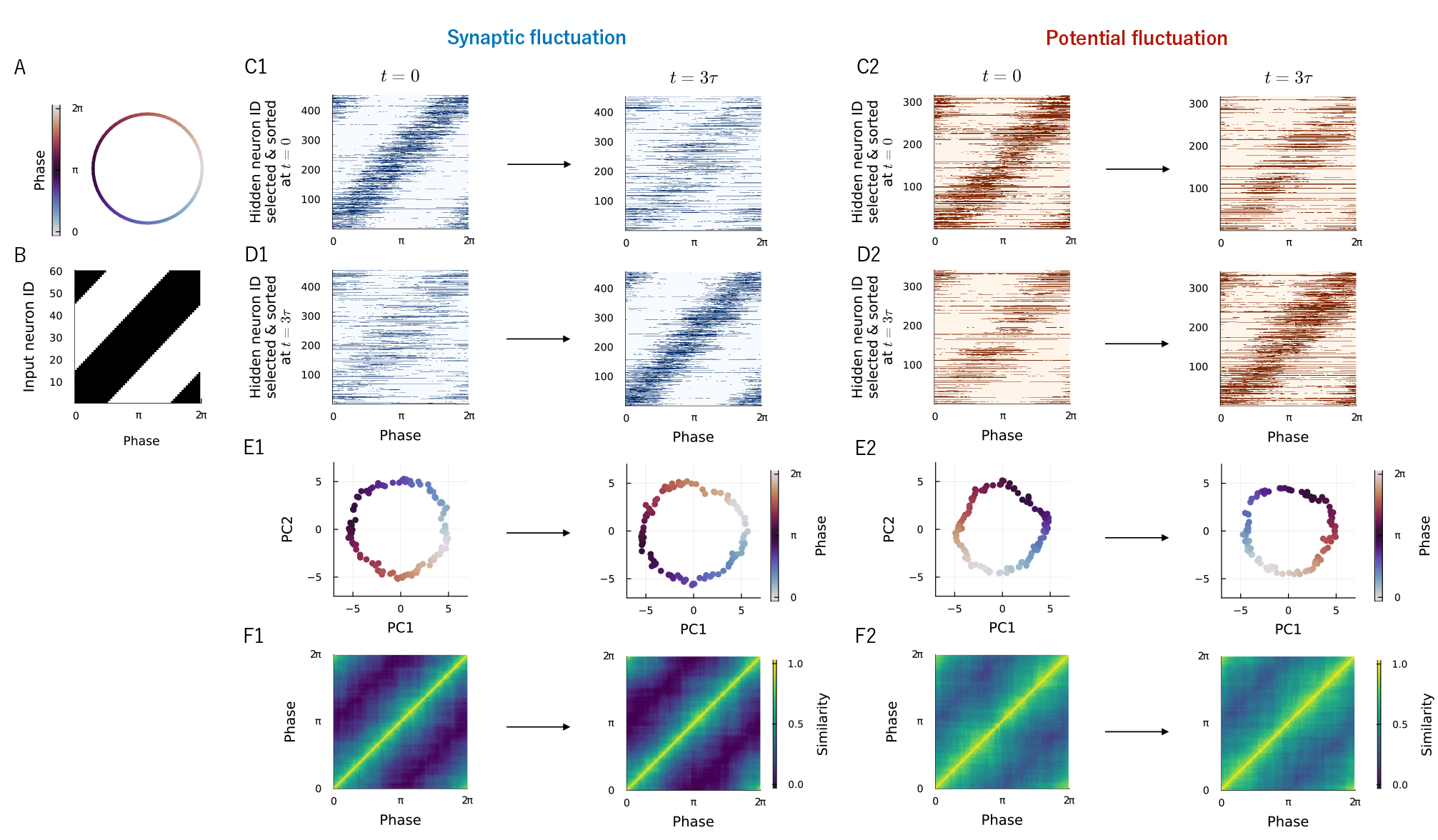
The structure of representations is preserved during drift under both mechanisms. (A) Stimuli organized in a ring structure. (B) Neurons in the input layer are tuned to specific phases of the ring structure and tile the entire phase as a population. (C-F) Representational drift from time *t* = 0 to 3*τ* for synaptic (C1-F1) and potential fluctuations (C2-F2). (C,D) Responses of subsets of hidden neurons, together representing the entire phase (horizontal axis) at *t* = 0 (C) and *t* = 3*τ* (D) respectively, sorted by neuronal tuning at corresponding time points. (E) PCA projections of stimulus representations show that both the mechanisms temporally preserve the ring structure (see Methods). (F) Cross-stimulus correlation analysis also shows that both mechanisms temporally preserve the representational similarity (see also Fig. S3). The strength of fluctuation *σ* was 1.0.

We then computed the transformed representations in the hidden layer and examined how they drifted under modest fluctuation strength (*σ* = 1.0, Fig. 2). We found that, under both fluctuation mechanisms, a subset of neurons together representing the entire phase lost its original activity pattern over time (Fig. 2C). However, another neuronal subset gained its activity in a compensatory manner to represent the entire phase (Fig. 2D). Experimental studies have also reported such phenomenon [11, 13]. Because of these compensatory changes in population activity, both mechanisms maintained the relationship between representations of different stimuli. Indeed, dimensionality reduction by principal component analysis (PCA) reproduced the ring structure at both the initial and the final time points (Fig. 2E). Likewise, the correlational structure between representations was preserved over time (Fig. 2F). These results indicate that the structure of representations is preserved across time during drift.

### Representational discriminability is maintained only under synaptic fluctuation with increasing drift magnitude

We next asked if the two fluctuation mechanisms would exert differential effects when the magnitude of drift is increased (Fig. 3). We indeed found differences in the discriminability of different stimuli. Under potential fluctuation, the correlation between representations became higher as the fluctuation strength was increased, even for a distant stimulus pair (Fig. 3A,B (right)). In contrast, synaptic fluctuation did not alter the representational correlation (Fig. 3A,B (left)). Representations in a lower dimensional space constructed by PCA showed an analogous trend: while potential fluctuation shrank the size of the ring, synaptic fluctuation did not even after larger drift (Fig. 3C,D). These results indicate that potential fluctuation impairs whereas synaptic fluctuation maintains the discriminability of stimuli as the fluctuation strength and drift magnitude increase. Notably, although potential fluctuation degrades discriminability with increasing strength, the resulting impaired structure remains stable over time (Fig. 3B,D and Fig. S4).

**Figure 3:**
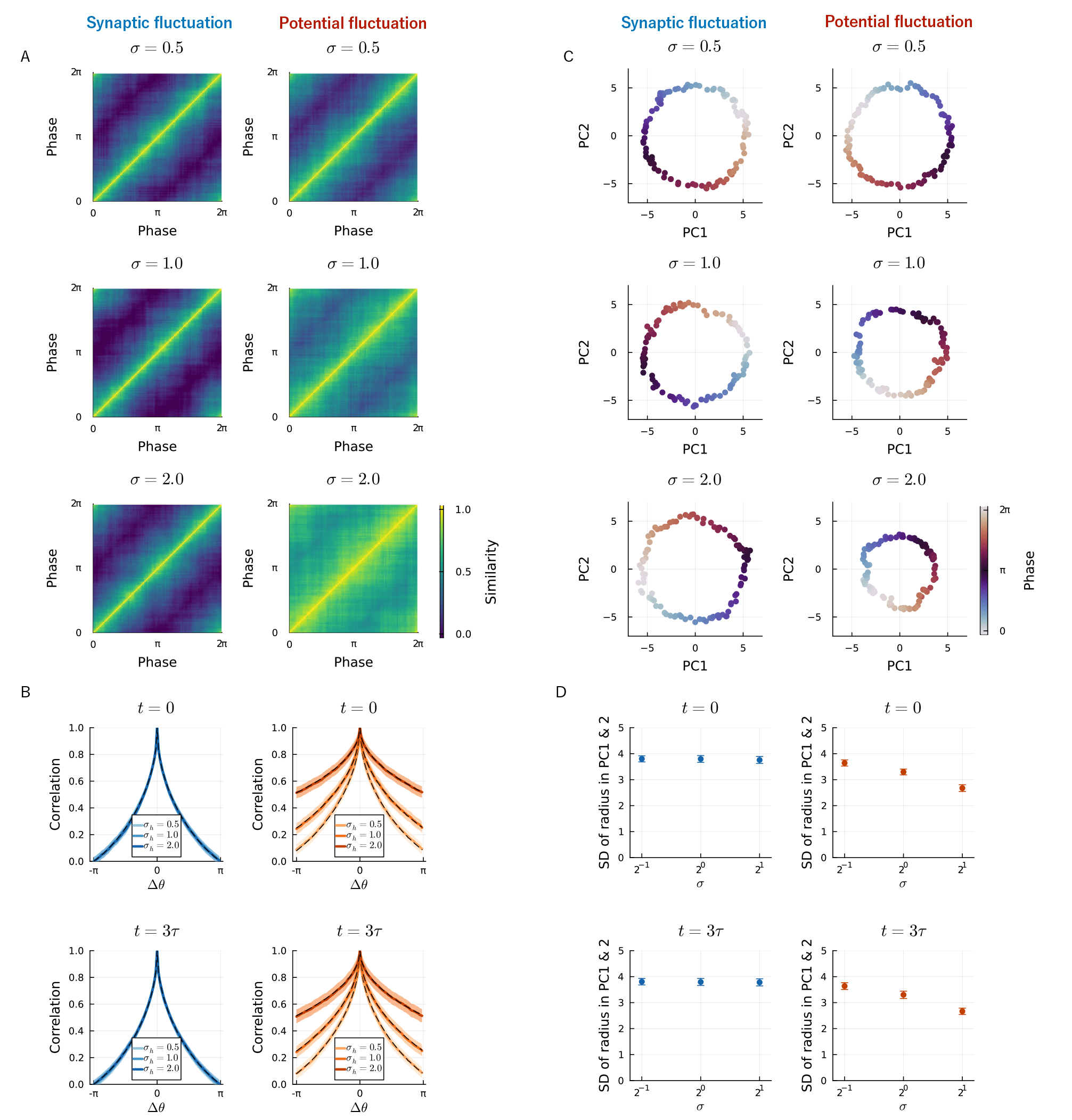
Synaptic but not potential fluctuations preserve representational structure under stronger drift. (A,B) Cross-stimulus correlation analyses show that potential fluctuations increase similarity across stimuli and reduce discriminability, while synaptic fluctuations preserve it. (C) PCA projections of stimulus representations for increasing fluctuation magnitudes (*σ* = 0.5, 1.0, 2.0). With potential fluctuations, the structure collapses as *σ* increases, whereas synaptic fluctuations maintain geometry (see Methods). (D) Standard deviation in PC space confirms this collapse under potential but not synaptic fluctuations. In panels (A) and (C), results shown are at *t* = 3*τ* (see Fig. S4 for consistent results at *t* = 0). Shaded regions (B) and error-bars (D) show variability across 100 simulations. Black dashed curves indicate theoretical predictions assuming infinitely large networks.

### Mechanisms underlying the stability and instability of representations during drift caused by two fluctuations

Thus far, we have observed that both synaptic and potential fluctuations preserve the structure of representations under moderate drift (Fig. 2), but exert differential effects under larger drift (Fig. 3). How do microscopic fluctuations translate into changes in the response of individual neurons that ultimately determine the population level stability? To address this issue, we visualized the responses of individual neurons in a selectivity space [41]. The selectivity space is spanned by neuronal responses to multiple stimuli, in which individual, hidden neurons are scattered according to their tuning (Fig. 4A). As we explain in detail below, combined with the equilibrium properties of synaptic and potential fluctuations (Fig. 4B), the dynamics of neurons within this space clarify the relationship between fluctuation mechanism, tuning of individual neurons, and emergent population properties.

**Figure 4:**
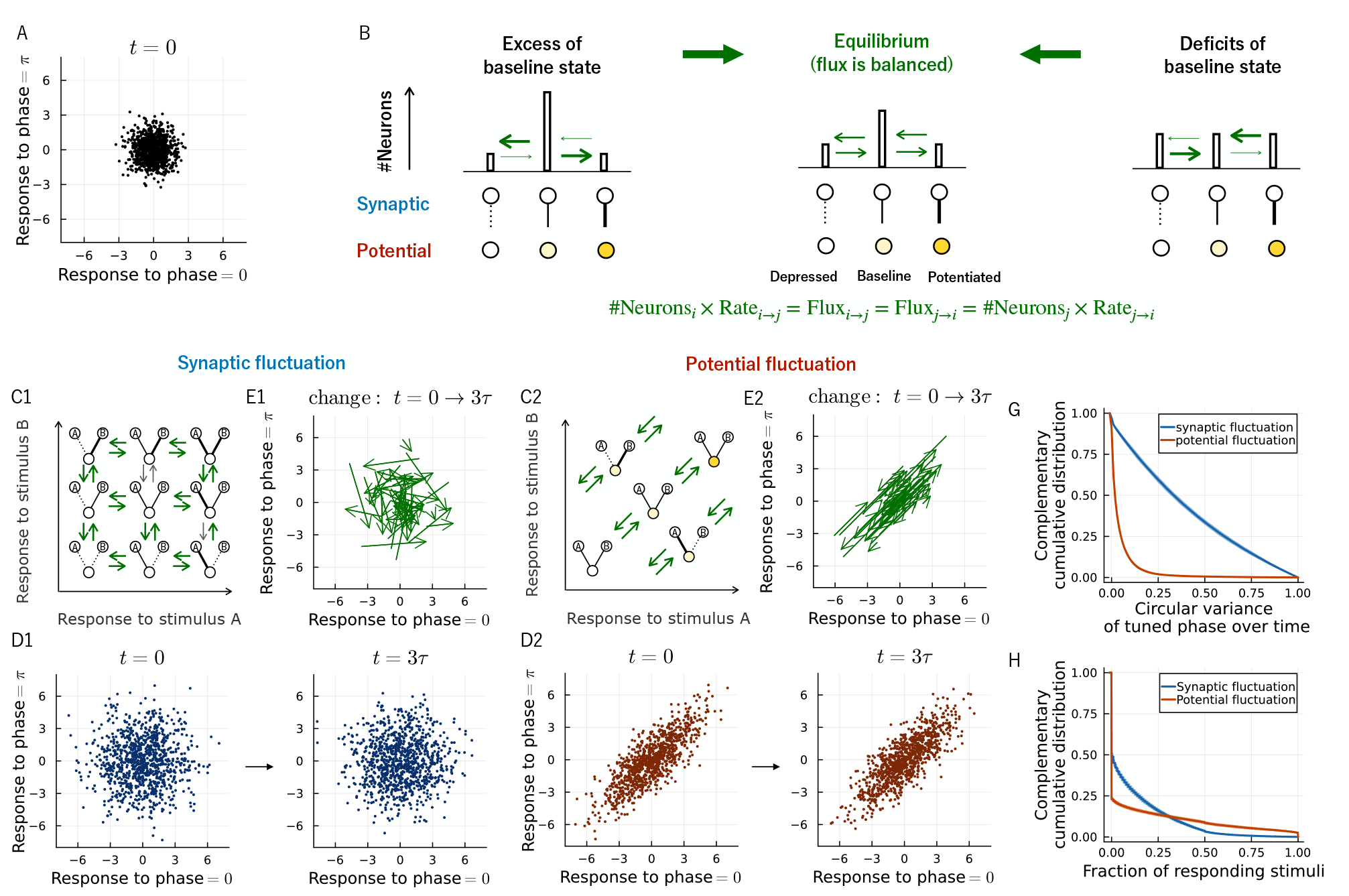
Mechanisms underlying the stability and instability of representations during drift caused by two fluctuations. (A) Selectivity space is spanned by neuronal responses to a set of stimuli. Each neuron’s tuning is represented as a point in this space. Input current is used as a proxy for response throughout this figure (see Methods). The example indicates the response to two stimuli with phase = 0, *π* (Fig. 2A,B), which produce non-overlapping representations at the input layer in the absence of fluctuation (*σ* = 0). (B) Schematic explaining the stability of distribution at equilibrium induced by synaptic and potential fluctuations. For simplicity of illustration, the synaptic weight and potential takes one of three states (depressed, baseline, or potentiated) and the number of neurons at each state is indicated by the height of bar. The middle panel shows the distribution of neurons at equilibrium where the inward flux into and the outward flux from the baseline are balanced. The left and right panels illustrate how perturbations to the equilibrium distribution spontaneously correct through homeostatic changes in fluxes: when the number of neurons in the baseline state is in excess (left) or in deficit (right), the resulting imbalance between inward and outward fluxes (green arrows) drives the system back toward equilibrium. (C) Schematics illustrating how individual neurons change their tuning under drift with synaptic (C1) and potential (C2) fluctuations. (D) Both mechanisms temporally preserve the distribution of neuronal tuning at a population level. (E) Tuning of individual neurons is significantly changed over time under synaptic fluctuations (E1) whereas constrained along the diagonal axis under potential fluctuations (E2). 5% of neurons are randomly chosen for visual clarity. (G) Distribution of variability in preferred phase during drift, measured by circular variance, shows differential tuning dynamics between the two mechanisms (see Methods). (H) Distribution of fraction of responding stimuli reveals the consequence of distinct tuning dynamics: potential fluctuation produces larger groups of neurons that respond to all the stimuli (= 1) or none of the stimuli (= 0) compared to synaptic fluctuation. Results at time *t* = 3*τ* are shown (see Fig. S5 for consistent results at *t* = 0). Curves and Shaded regions indicate mean and standard deviation over 100 simulations. The fluctuation strength is *σ* = 0 in panel (A) and *σ* = 2.0 in panels (D) through (H).

We first examined synaptic fluctuations using a simplified network in which each hidden neuron receives inputs selective to two stimuli (Fig. 4C1). Synapses stochastically transition between discrete strength states, allowing individual neurons to explore different response patterns over time. Neuronal responses reach an equilibrium in the selectivity space, even without interactions, as the fluctuation process homeostatically balances inward and outward fluxes for each state (Fig. 4B). Here, the flux represents the number of neurons transitioning from one state to another and is proportional to the number of neurons at the source of transitions. For example, a decrease in the number of neurons at the baseline state will reduce the outward flux from this state, but this imbalance will be spontaneously corrected by an increase in the inward flux due to the excess number of neurons in the remaining states. This equilibrium mechanism ensures that, although individual tuning changes substantially, the overall distribution of neurons in the selectivity space remains stable.

This logic described under the simplified scenario with two input neurons having discrete-valued weights can be directly applied to understand the behavior of the original model with many input neurons having continuous-valued weights. We found that the overall distribution of neurons in the selectivity space remains stable over time (Fig. 4D1) despite substantial change in tuning of individual neurons (Fig. 4E1). This corresponds to the preservation of representational structure across time described earlier (Fig. 2). Here, for clarity of visualization, we plotted the input current as a proxy for the neuronal response, which is sufficient for analyzing correlation properties captured on an ordinal scale (see Methods).

The effect of potential fluctuations can be understood similarly using fluxes in the selectivity space because synaptic and potential fluctuations exhibit identical dynamics (Fig. 4B) that shape the distributions of neurons at equilibrium (Fig. 4C2). Critically, potential fluctuation moves the neurons only along the diagonal direction (Fig. 4C2, E2) from the basal positions in the absence of fluctuation (Fig. 4A), keeping the preferred phases of neurons stable across time (Fig. 4G). Therefore, although the overall distribution of neurons remains stable over time (Fig. 4D2), its shape is more elongated obliquely under the presence of drift (compare Fig. 4D1 and 4D2). This gives rise to the higher correlation of representations under strong potential fluctuations described earlier (Fig. 3). This correlation also causes fraction of responding stimuli to concentrate more at 0 (no response to all stimuli) and 1 (response to all stimuli) compared to synaptic fluctuation (Fig. 4H).

In sum, temporal stability of representational structure emerges from equilibrium of neuronal tuning distribution despite drift, whereas shrinkage of representational structure emerges under strong potential fluctuation that causes excess positive correlation between neuronal responses to different stimuli.

### Both mechanisms improve reversal learning of single stimulus while only synaptic fluctuations improve reversal learning of multiple stimuli

We have seen that drift, under certain mechanisms and conditions, can preserve the structure of representations and overcome the perceived problem of stable stimulus discrimination. What, then, might be the beneficial role of representational drift? One hypothesis is that it facilitates associative, reversal learning, which involves not only the acquisition of new sensory associations but also the suppression or removal of old associations. Drift may facilitate the latter process as it breaks the old associations by changing the representations of sensory cues. To test this hypothesis, we utilized a reversal learning task where the valence of reinforcing signal associated with a sensory stimulus was switched at random timings. We used a linear readout trained by a Hebbian rule to report the valence of each stimulus (Fig. 5A,B). The timings of stimulus-reinforcer presentation as well as the rule reversal were determined following the Poisson process (Fig. 5C,D). Each time the network received a stimulus-reinforcer pair, the readout weights were updated according to the Hebbian associative plasticity [42] (Fig. 5B). Specifically, the readout weights were increased or weakly decreased if the corresponding hidden neurons were active or inactive at the moment of reward presentation. The readout weights were updated in an opposite manner for the presentation of punishment. We trained the readout for a certain period of time and observed its response to a stimulus immediately after the end of the training. The sign of the output of readout and the valence of stimulus should match if the readout weights were properly learned.

**Figure 5:**
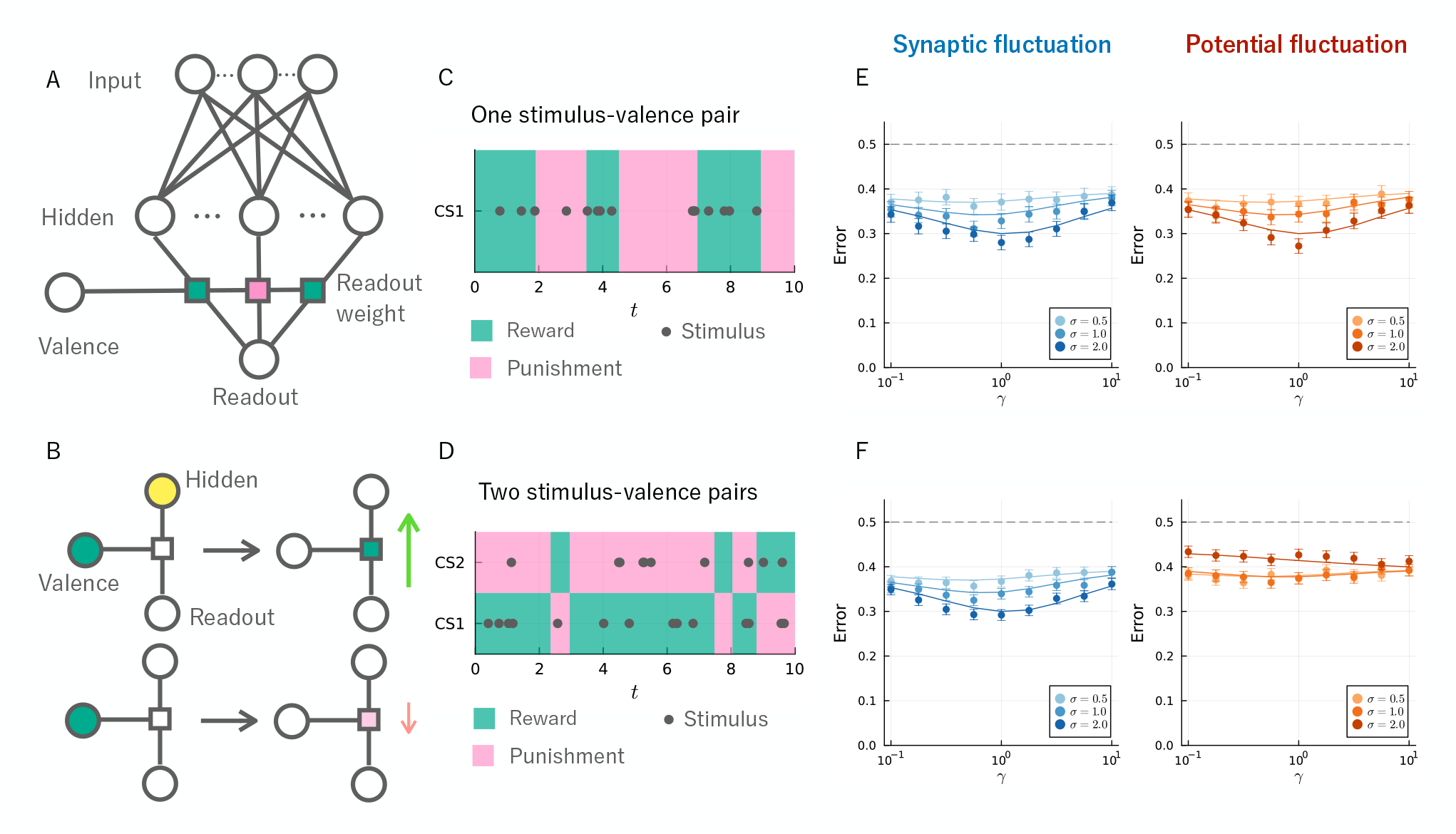
Beneficial effect of drift on associative, reversal learning. (A) Schematics of a modified model for associative learning. We added a linear readout integrating the activity of hidden neurons and another input neuron sending information about valence. The network learns to classify the stimuli to be rewarding (positive valence) or punitive (negative valence) by outputting positive or negative value. (B) Learning is implemented by Hebbian associative plasticity of readout weights. Upon receiving a reward, the readout weight is increased or weakly decreased if the corresponding hidden neuron is active (top) or inactive (bottom). (C) Sample trajectory of reversal learning signals. Application of stimulus-valence pair and reversal of valence occurs at random time points (see Methods). Readout weight is trained for 10 unit time and the classification accuracy of the readout was evaluated at the end of the training. (D) Sample trajectory of reversal learning signals for the case of learning two stimuli. Here, opposite valence signals are paired with the two stimuli, requiring discrimination of stimuli for learning. The classification accuracy was evaluated by observing the response of readout to stimulus 1 at the end of the training. (E,F) Dependence of classification error on drift parameters indicates that both mechanisms similarly improve reversal learning of single stimulus (E) while only synaptic fluctuation improves reversal learning of multiple stimuli (F). Dots and error bars indicate the mean and the bootstrapped standard deviation of 3000 independent simulations. Solid curves indicate theoretical predictions assuming infinitely large networks (Eqs. (7) and (13)).

We examined two scenarios: one requiring association of a single stimulus-valence pair (Fig. 5C), and the other requiring association of two stimuli with opposite valences where discrimination between stimuli is necessary (Fig. 5D). In the case of a single stimulus-valence pair, the classification error showed similar trends under both synaptic and potential fluctuations. The error decreased as *σ* increased and *γ* approached ∼ 1 (Fig. 5E). However, in the case of two stimulus-valence pairs, the classification error behaved differently between synaptic and potential fluctuations (Fig. 5F). While synaptic fluctuation reduced the classification error similarly to the single pair case, potential fluctuation increased the error when the magnitude of drift was large (*σ* = 2.0). This demonstrates that the effects of synaptic and potential fluctuations diverge under associative learning requiring discrimination.

The dependence of classification error on drift parameters can be understood intuitively. Large drift magnitude erases old associations by decorrelating current and past representations (Fig. 1E). The drift rate has a trade-off: faster drift helps discard old memories more quickly but also shortens the retention of recent ones (Fig. 1F). We confirmed analytically that learning is most efficient when the timescale of drift matches the timescale of valence switching (see Methods).

These results show that although drift with both mechanisms enables flexible updating of associations under a changing environment, synaptic fluctuation is superior because it retains discriminability even under large drift.

## Discussion

We have shown both similarities and differences in the effect of synaptic and potential fluctuations on representational drift and reversal learning. By aligning the magnitude and timescale of drift, we have identified qualitative differences that originate in the differences in fluctuation mechanisms. A crucial distinction lies in the way these mechanisms shape the relationship between multiple stimuli: whereas potential fluctuation at high magnitudes impairs stimulus discriminability by generating correlated representations, synaptic fluctuation maintains discriminability by keeping the structure of representations. Through an analysis in the selectivity space, we illustrated how these contrasting effects arise from different fluctuation mechanisms. Furthermore, examination of associative learning under the fluctuating environment revealed differential, functional consequences of these drift properties. When learning a single stimulus-valence pair where discrimination is unnecessary, both mechanisms enhanced associations under switching rules. However, when learning two stimulus-valence pairs, only synaptic fluctuation substantially improved the performance, as it induces significant drift without altering the representational structure necessary for accurate sensory discrimination.

Though we revealed the advantages of synaptic over potential fluctuations, potential fluctuations can be beneficial under certain situations. In contrast to synaptic fluctuations that substantially change neuronal tuning, potential fluctuations primarily modulate the overall responsiveness while keeping the preferred stimulus of neurons stable (Figs. 4D2,G). The latter meets the requirement of the neural null space hypothesis, which argues that the drift orthogonal to the readout can maintain stable downstream behavior [5]. Under potential fluctuation, neurons move along a single axis in selectivity space with small impact on neuronal tuning, generating drift-invariant, hidden neurons as a group (Fig. 4D2). If the readout weights of neurons within each group remain static, the output of readout would be invariant to drift because the distribution of neuronal tuning within each group is maintained under equilibrium. By confining the drift to such a low-dimensional space with small effects on the readout, potential fluctuation could confer stability to downstream behavior. On the other hand, we described that synaptic fluctuations are advantageous under the circumstance where readout weights can be tuned dynamically (Fig. 5). Therefore, synaptic and potential fluctuations have distinct beneficial effects depending on the stability of weights between hidden and downstream neurons. Recent experimental work has revealed that the passage of time drives changes in neuronal responsiveness (resembling potential fluctuation), while stimulus presentation drives changes in tuning properties (resembling synaptic fluctuation) [43]. Investigating the context in which differential fluctuation mechanisms are recruited presents an intriguing avenue for future research.

We have used the term “potential fluctuation” to make our result accessible by symbolically grounding our model in a biological context. However, we note that the implementation of potential fluctuation in our study need not to be confined to fluctuations of resting potential or spiking threshold mentioned in the model description. As shown in Fig. 4C-E, what distinguishes potential fluctuation from synaptic fluctuation is the concerted changes in neuronal responses to different stimuli. As long as this property holds, any biological process can implement the effect of potential fluctuation. For example, fluctuations in synaptic connections from input neurons that respond to all stimuli would have the same effect as the potential fluctuation described in this study.

We numerically illustrated the stability of relationships between representations during drift (Fig. 2), a notable phenomenon observed in multiple experiments [13, 16, 30]. While previous studies explained this stability based on the concept of equilibrium where the fluxes are spontaneously balanced [22, 30], these accounts lacked specific network mechanisms or numerical demonstrations, limiting their accessibility. Using the selectivity space, we complemented these studies by showing how synaptic fluctuation maintains the structure of representations despite substantial changes in the tuning of individual neurons (Fig. 4). An alternative mechanism for achieving stability has been proposed based on Hebbian and anti-Hebbian plasticity [24]. This approach differs from the mechanism assuming equilibrium because it achieves stability even with a few hidden neurons by incorporating plasticity in both feedforward and feedback connections (see Fig. S6 for the collapse under a few neurons with the equilibrium mechanism). Future work should address the general conditions required for stability, and we propose that the analysis in the selectivity space is useful to gain insight.

The differential effects of synaptic and potential fluctuations became apparent when the magnitude of drift was relatively large (*σ* ≥ 2) with autocorrelation approaching ∼ 0.1 (Fig. 1C). Experimental studies have reported various levels of drift with autocorrelation dropping to ∼ 0.2 [16, 43], although smaller values are yet to be observed perhaps due to factors such as the limitations in the duration of measurements. To evaluate the biological relevance of our findings, longer-term measurements will be required to examine the asymptotic behavior of drift dynamics.

We modeled fluctuation by the simple OU process for analytical tractability, but the qualitative results would generalize to wider class of fluctuations. The preservation of representational structure demands that the distribution of neurons in the selectivity spaces should be stable (Fig. 4), which is met if the fluctuation incorporates a homeostatic mechanism like the OU process. Furthermore, the increased representational correlation between different stimuli induced by potential fluctuation should be observed as long as the fluctuation changes the neuronal responses to every stimulus in a coordinated manner. Nevertheless, investigation of other types of fluctuations is important as quantitative detail of drift such as specific functional form between variance of potential fluctuation and discriminability of representations can depend on the details of fluctuations.

To identify the mechanisms and parameters underlying drift from experimental measurements, we need the information about the structure of representations in the upstream layer and the neuronal transfer function *ϕ* that specifies the relationship between the input current and the output spiking frequency [44]. High correlation between representations of different stimuli can result from either potential fluctuation or a large overlap in the representations in the upstream layer (Fig. 4), indicating the importance of characterizing the upstream activity to distinguish between these possibilities. In this respect, the *Drosophila* olfactory system represents a promising model because sensory representations have been measured comprehensively at two successive layers, with drift occurring in the postsynaptic layer [45, 46, 47]. It is also critical to examine the neuronal transfer function because it affects the relationship between the input current correlation and the activity correlation [44, 48, 49]. If drift originates in the input current dynamics, the transfer function can have a significant influence on the observed activity correlation. Given the challenge in characterizing the transfer function for all the neuronal types, there is a need to develop methods to address this uncertainty. One approach is to use mutual information instead of correlation to measure the representational similarity, as it is more robust to coordinate transformations [50].

In our model, we have simplified the biological complexity by omitting factors such as recurrent connections, activity-dependent plasticity, and spiking behavior for analytical tractability. Future work should build upon this foundation by incorporating more biological details to further investigate the drift mechanisms and their consequences.

## Method

### Random feedforward network with synaptic/potential fluctuation and correlation

#### Setting

We explain details of our drift model on a two-layer random feedforward network. We built our model based on previous work [42], largely following its notation and formulation. Input and hidden layer consists of *N*_*S*_ and *N*_*C*_ neurons. The activity of *i*-th neuron at time *t* in response to *n*-th stimulus is denoted as 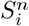 and 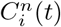 in input and hidden layers, respectively. We assumed that input layer representation is constant over time (i.e., does not drift). Neurons will be either active or inactive in response to stimuli, 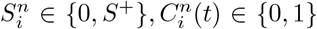, where *S*^+^ is a constant determined below. To simplify the analysis, we assumed that half of the input neurons respond to each stimulus and the response magnitude measured by Euclidean norm is the same for all the stimuli, which is 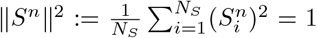. To satisfy this condition, a stimulus response of input neurons was set to 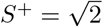. We denote a synaptic weight of feedforward connection from the *j*-th input neuron to the *i*-th hidden neuron as *J*_*ij*_. Under the absence of fluctuation, we assumed that the synaptic weights follow the Gaussian distribution 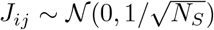 for simplicity and tractablity of the model. The hidden neurons are activated when the input current 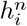 exceeds a certain threshold *T >* 0. Here, the input current was modeled by the weighted sum of input neurons’ activity:

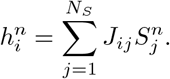

By using a step function *ϕ* (*ϕ*(*x*) = 1 for *x >* 0 and *ϕ*(*x*) = 0 otherwise), the activity of neurons in the hidden layer is expressed as

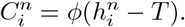

Threshold *T* was set such that the fraction of active hidden neuron 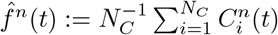 takes a predetermined value *f* ∈ [0, 1] at an initial time *t* = 0 for a stimulus *n* and was fixed thereafter. As described below, the value of 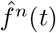 is approximately the same across stimuli *n* due to the constant magnitude assumption (∥*S*^*n*^∥ = 1) described above.

We introduced synaptic and potential fluctuations into the random feedforward network by adding two random variables 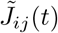 and 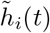 into the input current 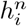 :

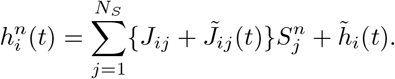

We assumed that these fluctuations proceed independently of the stimulus-driven neural activity, as observed experimentally and exploited in theoretical modeling [51, 7, 37, 38, 26]. Under this condition, the input current 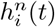 depends on time *t*. Each type of fluctuation was modeled by the following Ornstein-Uhlenbeck (OU) process:

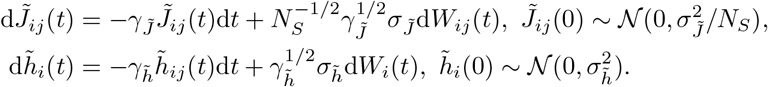

Here, 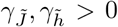 determines the timescale of fluctuation, 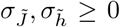 represents the amplitude of fluctuation, and *W*_*i*_(*t*), *W*_*ij*_(*t*) are independent standard Wiener processes such that 𝔼 [d*W*_*i*_(*t*)] = 0, 𝔼 [d*W*_*i*_(*t*)d*W*_*i*_(*t*^*′*^)] = 𝔼 [d*W*_*ij*_(*t*)d*W*_*ij*_(*t*^*′*^)] = *δ*_*ii*_*′ δ*_*jj*_*′ δ*(*t* − *t*^*′*^)d*t* where *δ*_*mn*_ and *δ*(*x*) are Kronecker delta and Dirac delta function. An explicit solution of the OU process implies that 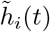 and 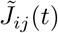 are normally distributed and have the mean 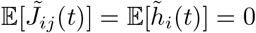 and covariance:

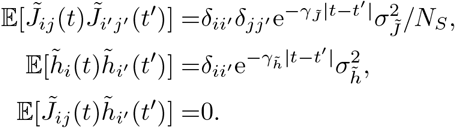

In this study, we focused on synaptic and potential fluctuations corresponding to 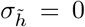 and 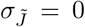 but analytical results below apply to general cases where both types of fluctuations simultaneously exist.

#### Statistical properties and their manifestations in a large network

Due to the normality of the input current 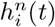, the statistical property of 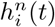 and its nonlinear transform 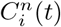 are fully characterized by the mean and the covariance of 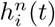. The input current has the mean 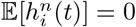 and the covariance:

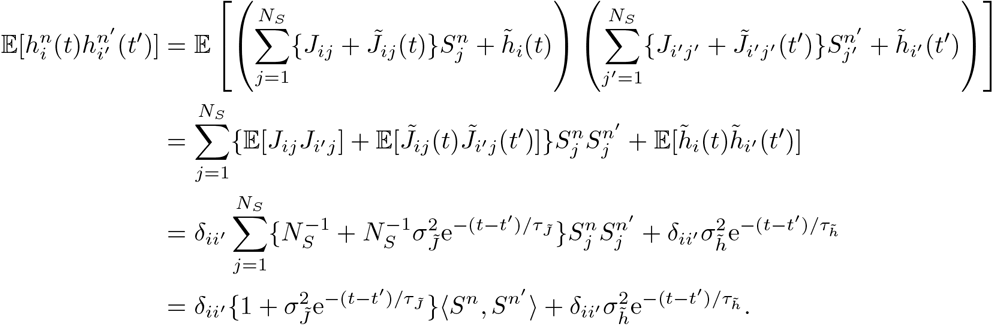

where 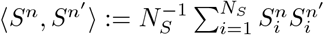 is the overlap between input layer representations. The variance is temporally constant, 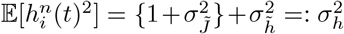, and is the same across stimuli due to the assumption ∥*S*^*n*^∥^2^ = 1. The variance *σ*_*h*_ determines mean activity depending on a given threshold *T* :

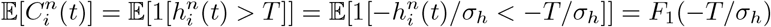

where 1[·] is an indicator function and *F*_1_ is a cumulative distribution function (CDF) of univariate standard normal distribution. We define a threshold 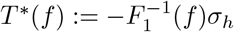 under which the mean activity takes *f* :

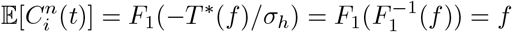

With this threshold value *T*^∗^(*f*), the correlation of input current across input *n* and time *t*

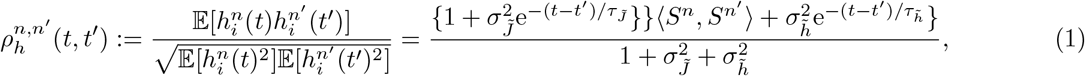

determines the activity correlation:

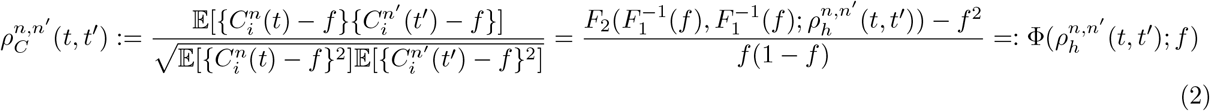

where *F*_2_(·, ·; *ρ*) is the CDF of bivariate standard normal distribution with correlation *ρ*. In Fig. S1(A), we plotted the relationship Φ between the correlation of the input current and that of the activity under *f* = 0.1. The activity correlation *ρ*_*C*_ drop rapidly than *ρ*_*h*_ near *ρ*_*h*_ = 1, reinforcing the idea that sparse representation (*f* = 0.1) improves the discriminability of similar stimuli [48].

These statistical properties determine the properties of representations when *N*_*C*_ is large enough to apply the law of large numbers (self-averaging). The active fraction of neurons at time *t* = 0 with an arbitrary threshold T is:

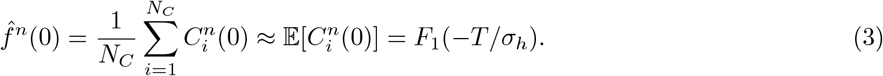

By substituting the threshold *T* = *T*_0_(*f*), which is set at *t* = 0 to satisfy 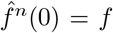, into the above formula, *T*_0_(*f*) is approximated by *T*^∗^(*f*) defined above:

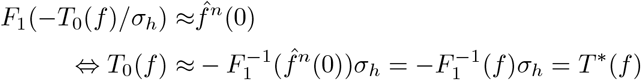

Then, the active fraction with the threshold *T*_0_(*f*) is approximately constant across time with the value *f* :

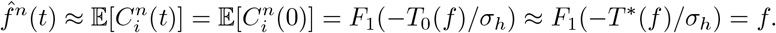

In the same way, the representational correlation approximately equals to statistical correlation 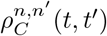 :

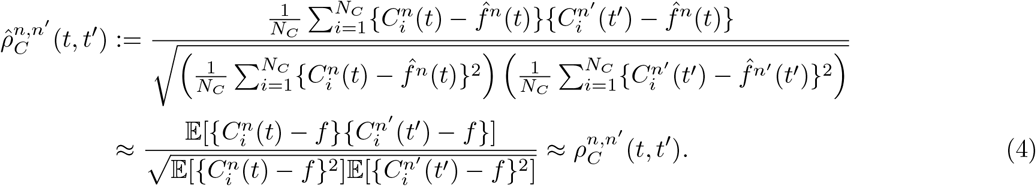

Therefore, the representational correlation 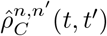, a key quantity in this study, can be characterized by 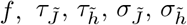 and 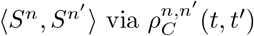 and 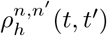 (Eqs. (2) and (1)).

#### Associative, reversal learning and optimal autocorrelation function

We describe the setting of associative, reversal learning and analytical expression of classification error by extending a previous work [42], where temporal dynamics were not considered in stimulus-valence association as well as neural representation.

#### Setting of associative, reversal learning

To implement associative learning, we added an input neuron to the network, sending information about valence denoted by *L*^*n*^(*t*) ∈ {±1} for each stimulus *n* at time *t* (Fig. 5(A)). By receiving pairs of valence signal *L*^*n*^(*t*) and neural response in the input layer *S*^*n*^(*t*), the network learns to classify the valence of stimuli solely from the input layer *S*^*n*^(*t*). The network outputs the result of classification *y*^*m*^(*t*) as a linear readout of representation in the hidden layer *C*^*n*^(*t*):

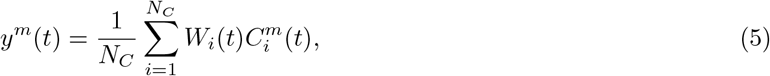

where *W*_*i*_(*t*) are readout weights. After learning, the linear readout is expected to take a positive/negative value in response to a stimulus with positive/negative valence (Fig. 5(A)). To implement this, readout weights *W*_*i*_(*t*) were learned by the associative Hebbian rule every time the network received a stimulus-valence pair (Fig. 5(B)):

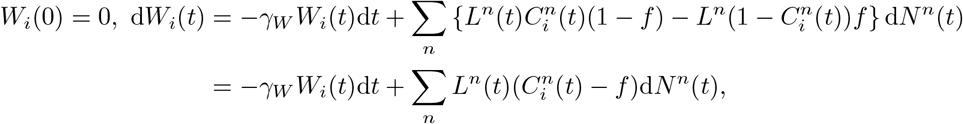

which is solved as:

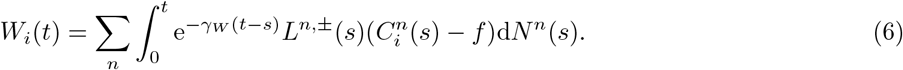

Here, *γ*_*W*_ represents a spontaneous decay rate of readout weight, which was introduced just for avoiding divergence, and 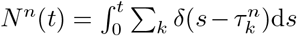 indicates the number of *n*-th stimulus-valence pair up to time *t* where 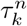 indicates the time of *k*-th occurrence of a *n*-th stimulus-valence pair. This equation indicates that when positive [negative] valence is received, readout weights *W*_*i*_(*t*) are updated in the direction of valence (1 − *f*) [−(1 − *f*)] if corresponding hidden neuron is active 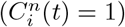 and weakly updated in the opposite direction by −*f* [*f*] if it is inactive 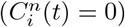.

We assumed that the timing of occurrence and the rule reversal of stimulus-valence pairs is unpredictable, which would be plausible in natural environment, but follows a known specific timescale, information that the network possibly acquired through longer-term learning or evolution. To model such conditions, we used the Poisson process *N* ^*n*^(*t*) = 0, 1, 2, … with rate *λ* [unit time^−1^] to set the timing of stimulus-valence pairs and the continuous time Markov chain *X*(*t*) ∈ ±1 with uniform initial probability ℙ (*X*(0) = ±1) = 1*/*2 and symmetric transition rate *r* = 1*/*2 [unit time^−1^] to set the timing of rule reversal (Fig. 5(B)). Here, we set a unit of time by the timescale of reversal *τ*_*X*_ = (2*r*)^−1^ without losing generality (note that the autocorrelation of *X* is 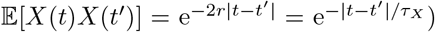 . Based on these processes, we instantiated two situations to illustrate the difference between synaptic and potential fluctuations. The first is learning a single stimulus-valence pair where discrimination is not required (Fig. 5(B) (top)). Valence *L*^1^(*t*) was set by *L*^1^(*t*) = *X*(*t*). The other is learning two stimulus-valence pairs with opposite valence (Fig. 5(B) (bottom)). We assumed that there is no overlap between representations in the input layer ⟨*S*^1^, *S*^2^⟩ = 0 and that the two stimuli were assigned with valence with opposite signs *L*^1^(*t*) = −*L*^2^(*t*) = *X*(*t*). This setting enhances the difference between synaptic and potential fluctuations.

The network was trained for a certain period [0, *t*_end_) following Eq. (6) and the classification error was tested by applying a single stimulus at the end of the training. The classification is correct if the signs of readout *y*^1^(*t*_end_) and valence *L*^1^(*t*_end_) are the same and wrong otherwise. The result of classification is equivalently expressed by a summary variable *g*^*m*^(*t*) := *y*^*m*^(*t*)*L*^*m*^(*t*), indicating correct/wrong by a positive/negative value. The test time point was set as 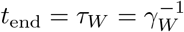, the longest autocorrelation time among the processes because *W* was assumed to have slowest decorrelation to minimize its effect on learning. With this setting, the result of classification is expected to approach a stationary value.

#### Analytical expression of a classification error

We derive the analytical approximation of the classification error as follows. We first empirically observed that *g*^*m*^(*t*) distributes normally (Fig. S7), implying that the classification error can be approximated by the signal-to-noise ratio (SNR) of *g*^*m*^(*t*):

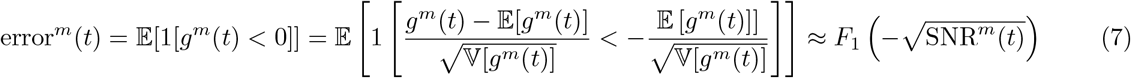

where SNR^*m*^(*t*) := 𝔼 [*g*^*m*^(*t*)]^2^*/*𝕍 [*g*^*m*^(*t*)].

To calculate the mean and the variance of *g*^*m*^(*t*), we substitute Eqs. (5) and (6) to the definition *g*^*m*^(*t*) = *y*^*m*^(*t*)*L*^*m*^(*t*):

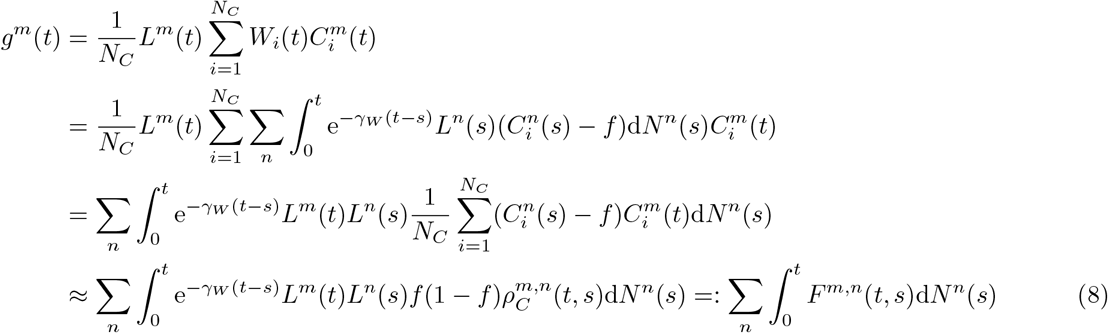

Where 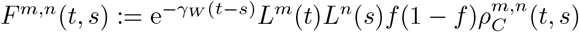. The mean of *g*^*m*^(*t*) is:

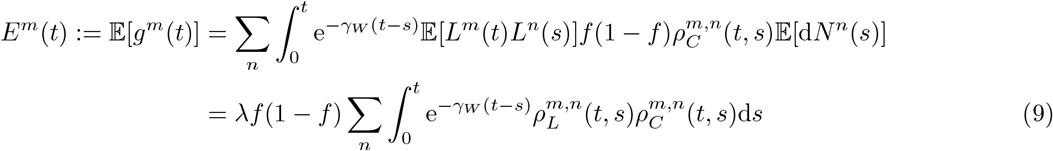

where 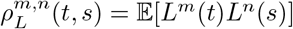 is the correlation between *L*^*m*^(*t*) and *L*^*n*^(*s*). The variance of *g*^*m*^(*t*) is decomposed as follows:

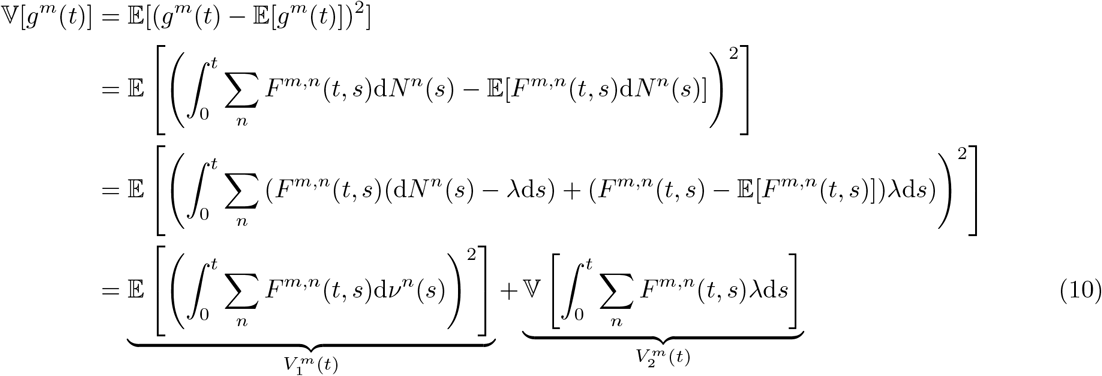

where d*ν*^*n*^(*s*) := d*N* ^*n*^(*s*) − *λ*d*s*. Cross-term vanishes due to 𝔼 [d*ν*^*n*^(*s*)] = 0 as well as the independence between the signal arrival 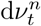 at time *t* and the state of network and valence *F* ^*m,n*^(*t, s*) just before time *t*. We calculate the first term by noting that 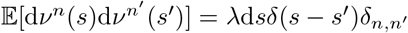 :

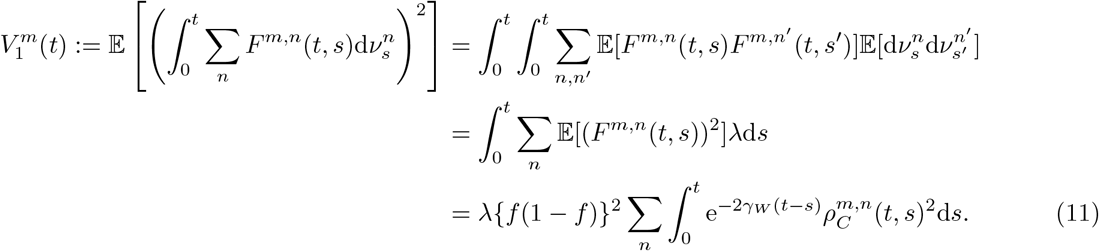

We expand the second term as follows:

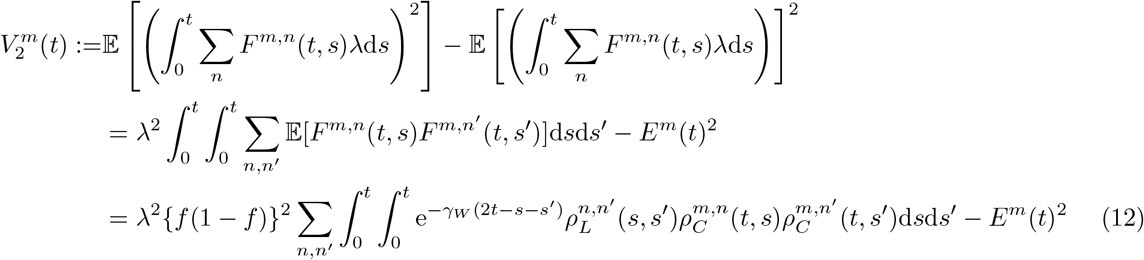

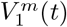 and 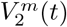 represent the variance originating in the random timing of stimuli-valence occurrence and reversal, respectively. We can now represent the classification error by substituting the following SNR into Eq. (7):

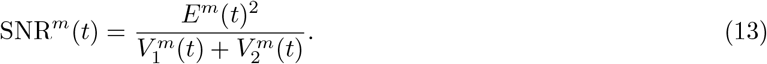

with 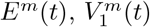, and 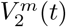 given by Eqs. (9), (11), and (12).

#### Optimal drift property for minimizing the classification error

We discuss the optimal drift property 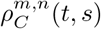 that minimizes the classification error given by Eqs. (7) and (13). Given that the error is a decreasing function of SNR (Eq. (7)), this optimization problem is identical to the maximization problem of the SNR (Eq. (13)). We derive the necessary condition of the optimal drift by taking the functional derivative of SNR^*m*^(*t*) with respect to 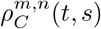 . To this end, we take the derivative of 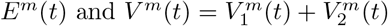:

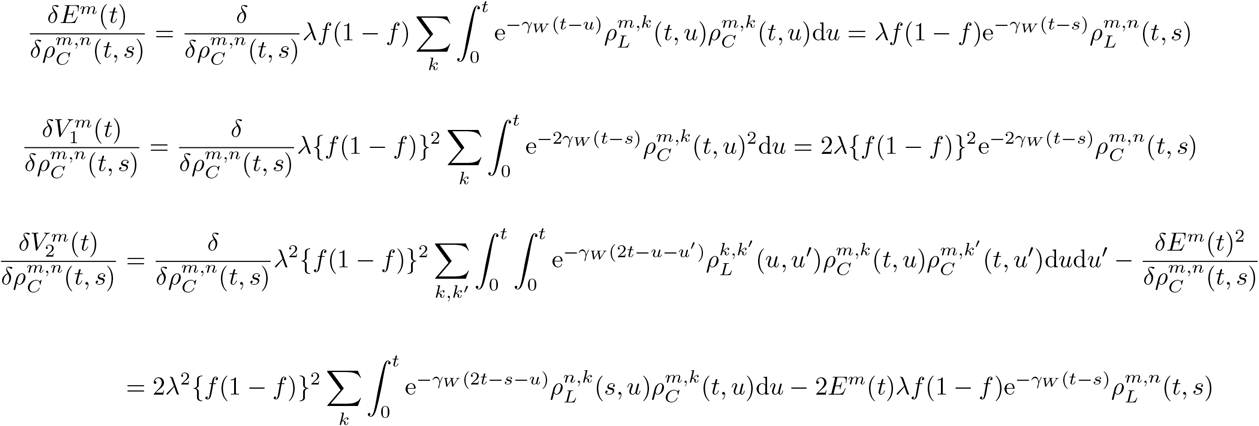

By setting 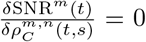 and substituting the above functional derivatives, we obtain the following stationary condition up to the multiplicative constant independent of *n* and *s*:

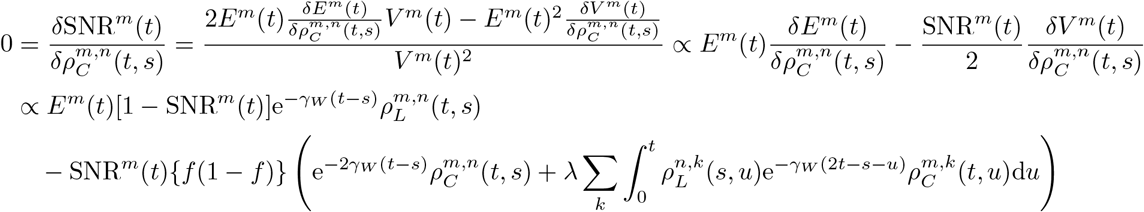

which implies that the optimal drift 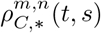 should satisfy

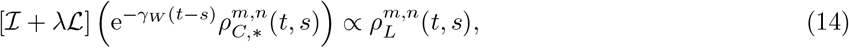

where 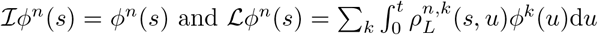 are the linear operators, and the proportionality constant is determined with the boundary condition 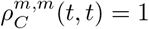.

To gain an intuition on the optimal drift property indicated by Eq. (14), we examine the regime where *λ* is small, which allows some analytical progress. When *λ* is small enough so that 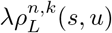, which is originated in 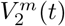, is negligible, we explicitly obtain the solution of Eq. (14):

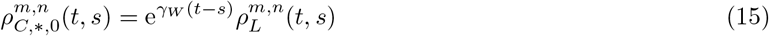

This result is also reproduced by using the Cauchy-Schwarz inequality when ignoring 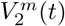. Indeed, when 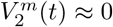, the SNR (*E*^*m*^(*t*)) is bound by a 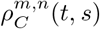 -independent quantity:

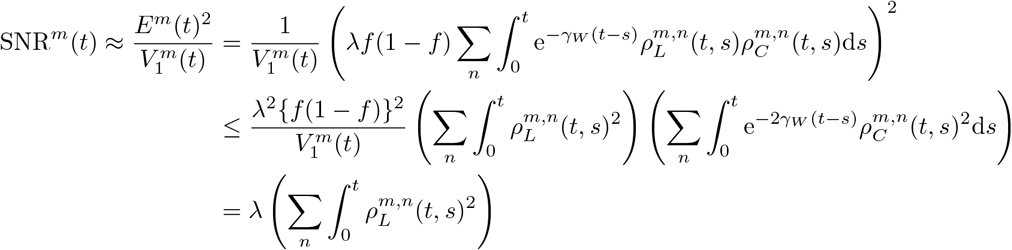

where the equality holds when and only when 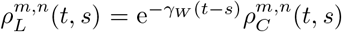 for all the stimulus *n* and time *s* ∈ [0, *t*]. When the decay of readout weight is negligible (*γ*_*W*_ ≪ 1 and e^−*γ W* (*t*−*s*);^ ≈ 1), Eq. (15) indicates that the optimal strategy is to match the drift dynamics 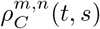 to the reversal dynamics of valence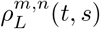. The intuition here is that the correlation between the current representation 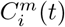 and the past representation 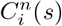 contributes to the signal *E*^*m*^(*t*) only when the valence *L*^*n*^(*s*) having been associated with 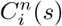 at past is informative for inferring the current valence *L*^*m*^(*t*), but contributes to the noise 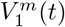 irrespective of whether the valence associated at past is informative or not. The result is also analogous to the Bayesian and information theoretical arguments that the optimal information processing requires an internal model that statistically predicts and emulates the environment.

To see the robustness of the zero-th order approximation (Eq. (15)), we examine the first order perturbation 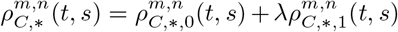 by regarding *λ* as a small parameter. The optimality condition (Eq. (14)) in the first order of *λ* is:

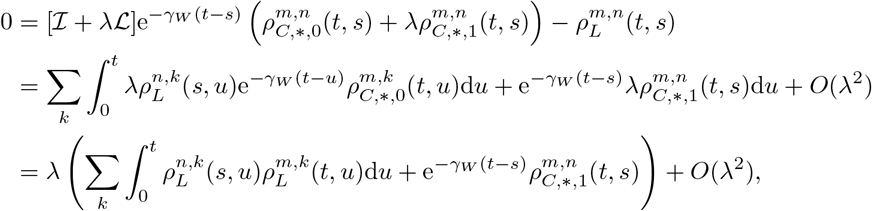

which implies

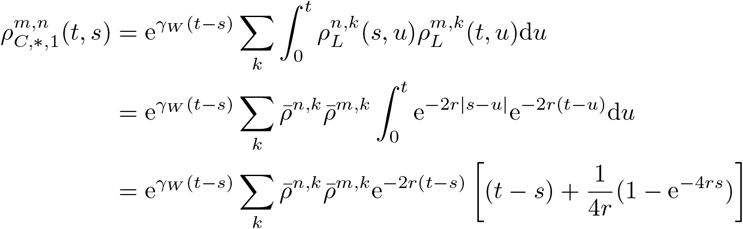

where we assume 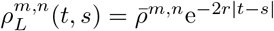 in the second line to examine the temporal and stimulus dependence individually. Although there is an additional multiplicative factor, the first order perturbation *ρ*_*C*,∗,1_ depends on the time primarily through e^−2*r*(*t*−*s*)^ as does the zero-th order solution 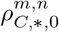 depends on it through 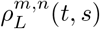. The dependence on stimulus *n* would also remain similar because if stimuli *m* and *n* has a high positive correlation 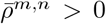 *>* 0, they would have similar correlations to other stimulus *k* so that 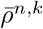 and 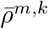 tends to have the same sign, i.e., 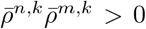, and the same discussion applies for negative correlation 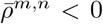 . Numerical solution also supported that the optimal time course of drift is robust to the increase of *λ* (Fig. S8(A)). A more complete discussion would be possible by noting that Eq. (14) has a relevant form with Fredholm integral equation of the second kind so that we may obtain more explicit solution as Neumann series when *λ* is not so large. It remains to be investigated to what extent the result is robust to increases of *λ* and whether qualitatively different behaviors emerge above certain *λ* (Eq. (15)).

We next describe how the derived optimal drift relates to the result of simulation in the reversal learning, assuming that the zero-th order solution (Eq. (15)) well captures the optimal drift. In our simulation, 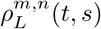 takes the following form depending on the situation: (1) 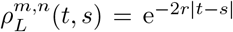 under the presence of a single stimulus-valence pair, and (2)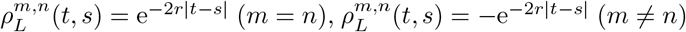 [under the presence of two stimulus-valence pairs with opposite valence. In either situation, the temporal correlation exponentially decays. To realize such decay in drifting representations, Fig. 1 (C) as well as Eq. (1) suggests that the amplitude of fluctuation 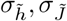 should be sufficiently large. Otherwise, there remains a substantial autocorrelation even after a long time. This explains why increasing 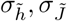 decreases the classification error in reversal learning (Fig. 5(D)). In the second situation, the optimal performance demands the correlation between representations of two stimuli to be negative. Under our setting, the minimum possible correlation between two representations is zero (uncorrelated), which is attained by synaptic fluctuation when there is no overlap in input representations. It would be interesting to consider the network mechanism that introduces negative correlation in representations, such as anti-Hebbian plasticity, depending on the negative correlation in valence signals.

#### Finite size effect

To complement the analysis, we show analytical results for the classification error under the moderate size of *N*_*C*_. Because the relationship between the classification error and the SNR does not change, we modify the calculation of mean and variance of *g*^*m*^(*t*).

Although we had used the law of large numbers to approximate the sum over hidden neurons 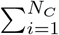, here we use the central limit theorem to evaluate the variance as well:

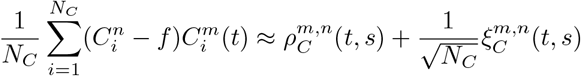

Where 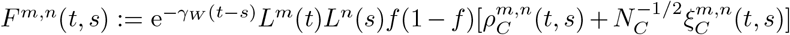 . We used the central limit theorem in the last line and defined a zero mean Gaussian random variable 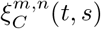 with the following covariance

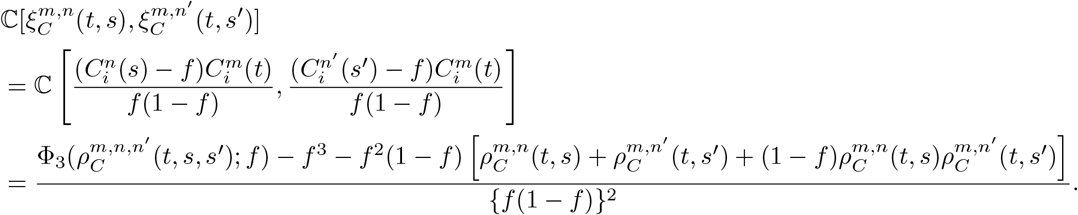

In the last line, we defined 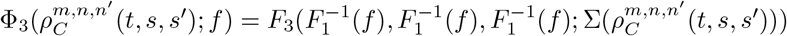 where *F*_3_(·, ·, ·; Σ) is the CDF of trivariate normal distribution with the covariance matrix Σ, and 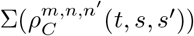 is a covariance matrix with three non-diagonal elements 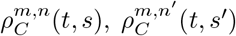 and 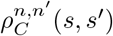 . In numerical calculation, we computed *F* ^3^ by the Monte-Carlo approximation with 10000 samples.

With this modification, *g*^*m*^(*t*) is expressed by replacing *F* ^*m,n*^(*t, s*) in Eq. (8) with 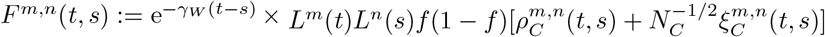. The mean of *g*^*m*^(*t*) reproduces the original expression:

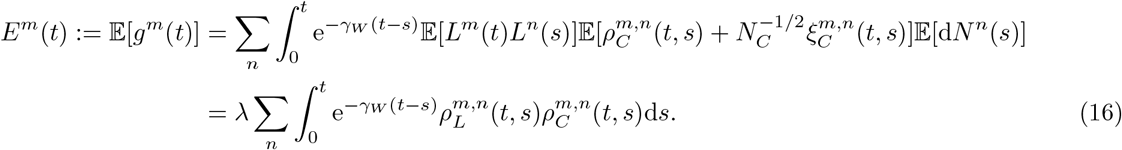

The decomposition of variance in Eq. (10) also holds. The first and second terms of variance are slightly modified as follows:

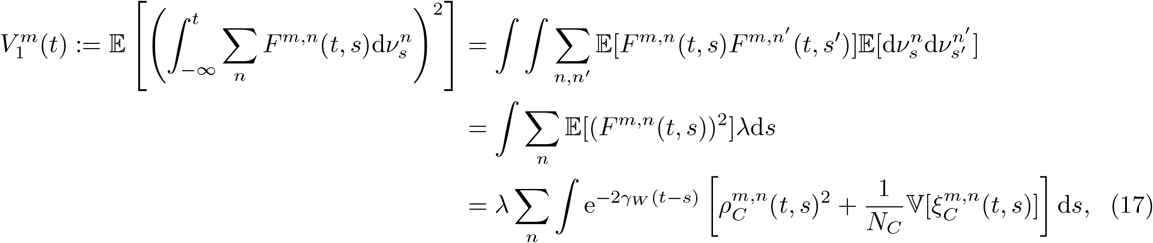

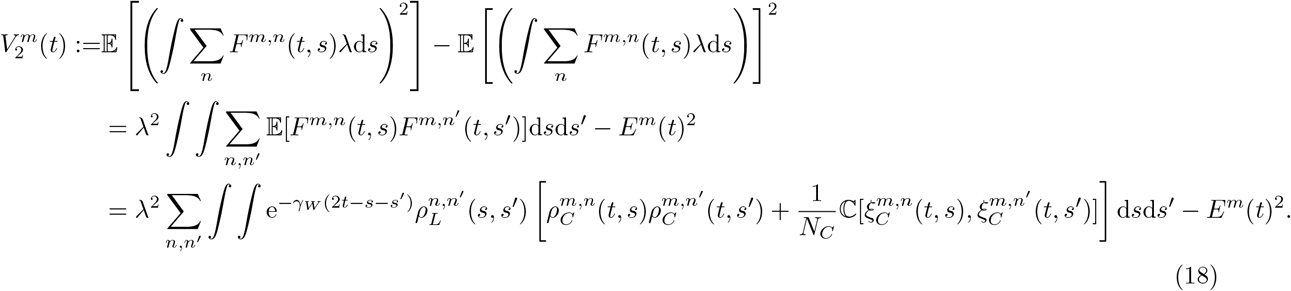

With these modifications, the classification error is represented by substituting the modified formula of 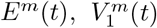, and 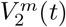 (Eqs. (16), (17), and (18)) into Eqs. (4) and (13). The validity of this approximation was confirmed by numerical simulation with *N*_*C*_ decreased from 1000 to 100 (Fig. S8(B)). The finite size effect was negligible at *N*_*C*_ = 1000 but significant at *N*_*C*_ = 100.

## Details of simulation

### Network parameters

Throughout this manuscript, we set *N*_*S*_ = 100, *N*_*C*_ = 1000, and *f* = 0.1 unless otherwise indicated. In Figure 5, we additionally set *λ* = 1.0 [unit time^−1^] and *r* = 1.0 [unit time^−1^].

### Stochastic processes

The Poisson process and the Markov chain used for the reversal learning were simulated by the direct method (see Eqs. (10a) and (10b) of Ref. [52]). The OU process used to introduce the synaptic and potential fluctuations was simulated by using the exact updating formula valid for arbitrary time steps (see for example Eq. (1.10) of Ref. [53]). The time step was determined by the direct method in the reversal learning (Fig. 5) and was fixed to 0.1 unit time in all the other simulations.

### Model and analysis in Figs. 2-4

*P* stimuli were associated with phase variables 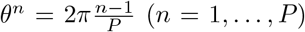 to express the ring structure. To reflect this structure, the representation at input layer was modeled such that each input neuron responds to half of the phases with a unique starting point: 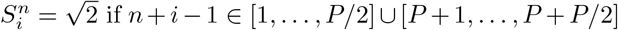. Here, *P* was taken to be an even number and the condition 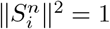 was satisfied.

In the analysis of temporal variability of preferred phase (Fig. 4G), time points where 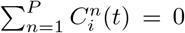 or 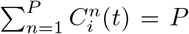 were omitted because the response is uniform across phases and therefore the preferred phase cannot be defined. The preferred phase for each cell at each time point was calculated as 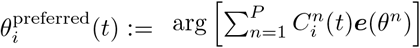, where ***e***(*θ*) is a unit vector with direction *θ* and arg[***v***] represents the direction of vector ***v***. The temporal variability was calculated using circular variance from directional statistics: 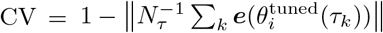 where *τ*_*k*_ (*k* = 1, …, *N*_*τ*_) are time points where the preferred phase is defined and *N*_*τ*_ is the number of such time points.

### Notes on the scale of PCA projection (Figs. 2 and 3)

PCA projections were calculated independently for each time point and drift parameter. Importantly, the scales of the principal components are comparable across different time points and drift parameters because PCA is an orthogonal transformation that preserves the Euclidean norm of the original neuronal response space.

### Visualization in selectivity space (Fig. 4)

For the purpose of visualization, we plotted the input current 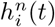 as a proxy for the neuronal response 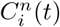 because it better reflects the actual correlation due to the normality of 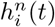 . Although this normality stems from the assumption of normality of synaptic weights in our model, it is expected to hold under broad situations as 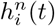 reflects an approximate linear sum of numerous independent or weakly correlated inputs under which the central limit theorem confers normality [54, 55, 56]. We further note that the correlation in input current 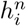 and activity 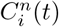 has a monotonic, albeit nonlinear, relationship (Fig. S1A; see also Ref. [57]), ensuring that our discussion based on 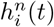 applies equally to 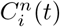 for correlation properties captured on an ordinal scale.

### Intuitive explanation of why the fraction of active neurons remained constant

We provide intuitive explanation of why the fraction of active hidden neurons remained constant without adaptively tuning the threshold *T* (Fig. 1). In a random, feedforward network, the input current to each neuron can be regarded as a sample from a certain distribution set by network statistics. As the number of hidden neurons *N*_*C*_ increases, the fraction of input current exceeding threshold *T* converges to a tail probability of a distribution by the law of large numbers. Because the fluctuation of our model preserves the variance over time, the tail probability remains constant, and so does the fraction of active neurons (see Eq. (3)). The agreement with analytical results indicates that *N*_*C*_ = 1000 is sufficient for applying the law of large numbers. Note that when calculations are restricted to responsive neurons, autocorrelation may be underestimated and the fraction of active neurons may be overestimated (Fig. S2).

## Acknowledgment

We thank Kensuke Yoshida, Genki Shimizu for helpful advices; Toshitake Asabuki, and Kazama lab members for valuable feedback on manuscript. K.N. was supported by RIKEN Special Postdoctoral Felloship and a grant from Japan Science and Technology Agency (ACT-X grant number JPMJAX24LC). This work was supported by a grant from RIKEN, Toray (23–6402), and Japan Science and Technology Agency (CREST grant number JP24028095) to H.K.

## Data availability

All code used in this paper are deposited at GitHub (https://github.com/rotala17/drift-feedforward). Simulation data that required extensive computation time are also deposited in the repository.

## Author contributions

K.N., K.E., and H.K. designed the research, K.N. performed the analyses. K.N. and H.K. wrote the paper swith input from K.E.

---

**Figure S1:**
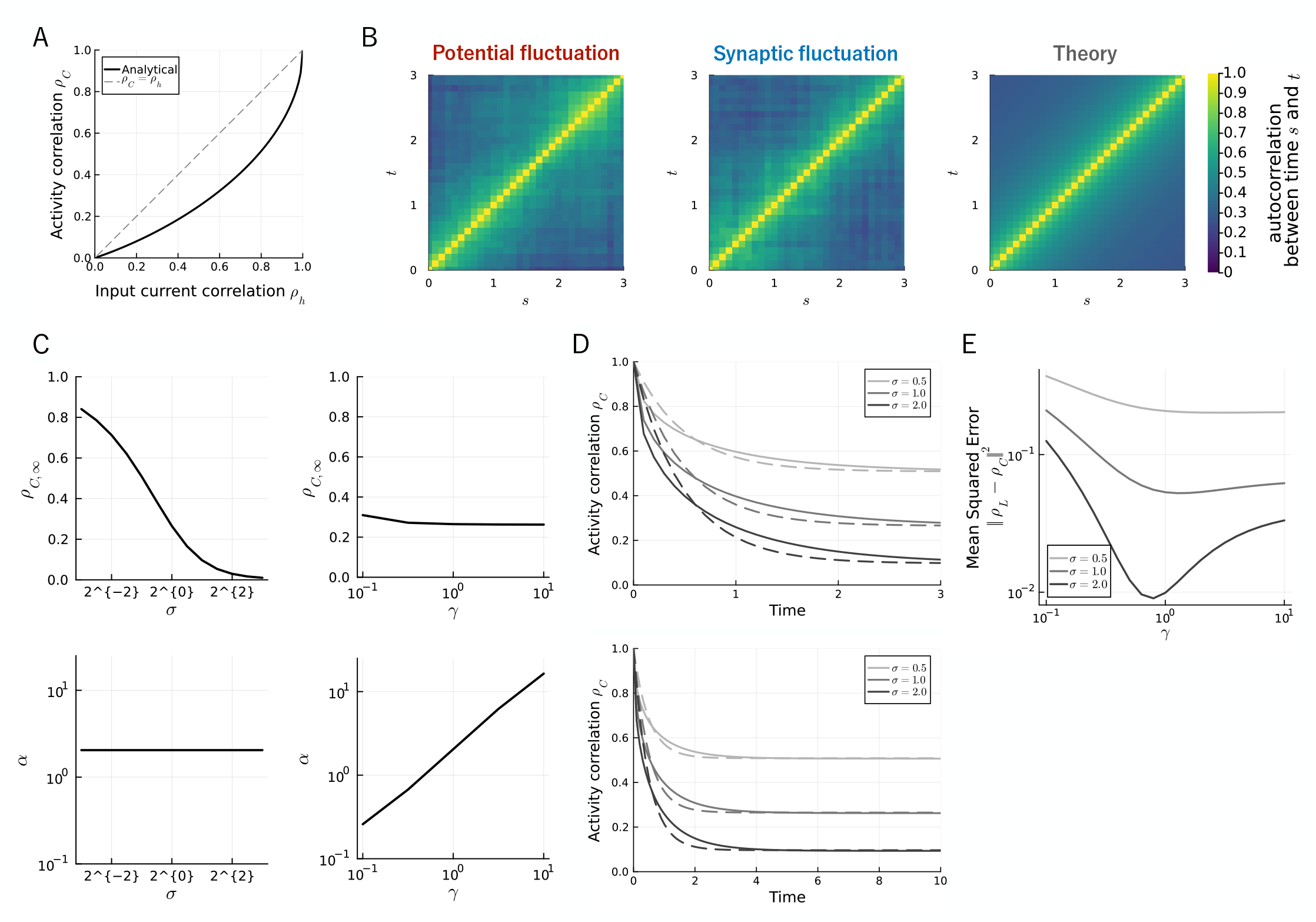
Autocorrelation of hidden layer representations. (A) Relationship between autocorrelation of input current *h* and activity *C*. Solid curve indicates the analytical expression (Eq. (2)). Dashed curve indicates the line *ρ*_*C*_ = *ρ*_*h*_. (B) Auto correlation of hidden layer representations between time *s* and *t* for synaptic fluctuation (left), potential fluctuation (middle), and analytical expression (Eq. (4)) (right). (C) Characterization of representational autocorrelation 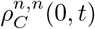 by an exponential decay function *ρ*_exp_(*t*) = (1 − *ρ*_∞_) exp(−*αt*) + *ρ*_∞_. We fit parameters of the function to the autocorrelation function (Eq. (4)) under the fixed drift parameters *σ, γ* and plot the relationship between drift parameters *σ* (left) and *γ* (right) and fitted parameters *ρ*_∞_ (top) and *α* (bottom). (D) Comparison between the representational autocorrelation 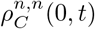 (solid line) and the exponential decay function *ρ*_exp_(*t*) with fitted parameters (dashed line). We indicate the autocorrelation over *t* ∈ [0, 3] (top) and *t* ∈ [0, 10] (bottom). Obtained parameter values are (*ρ*_∞_, *α*) = (0.51, 2.0), (0.27, 2.0), and (0.10, 2.0) for *σ* = 0.5, 1.0, and 2.0, respectively. (E) *γ*- and *σ*-dependence of squared error between the autocorrelation of representation 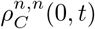 and that of valence 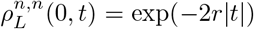 with *r* = 0.5. When fitting parameters in panels (C,D) and calculate the error in panel (E), we used the square error 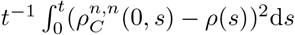 with *t* = 10 where *ρ*(*t*) = *ρ*_exp_(*t*) in panels (C,D) and 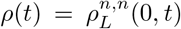 in panel (E). Network parameters were *N*_*C*_ = 1000, *N*_*S*_ = 100, *f* = 0.1, *σ* = 1.0, and *γ* = 1.0, unless otherwise indicated in each panel.

**Figure S2:**
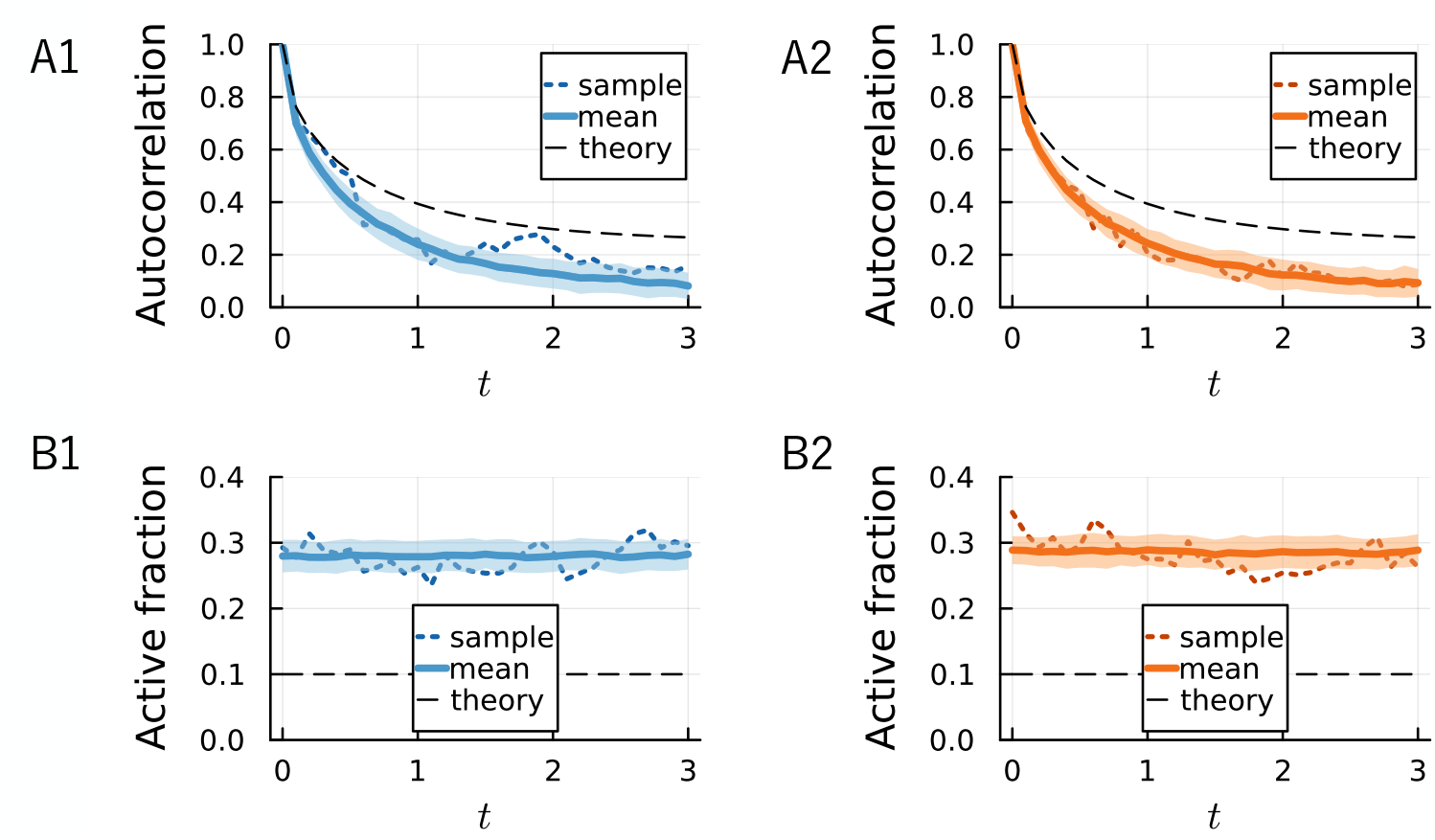
Effect of selecting responsive neurons. (A) Autocorrelation of drifting representation over time calculated based on the hidden neurons chosen in Figure 1B under potential (A1) and synaptic (A2) fluctuations. (B) Fraction of active neurons over time calculated based on the hidden neurons chosen in Figure 1B under potential (A1) and synaptic (A2) fluctuations.

**Figure S3:**
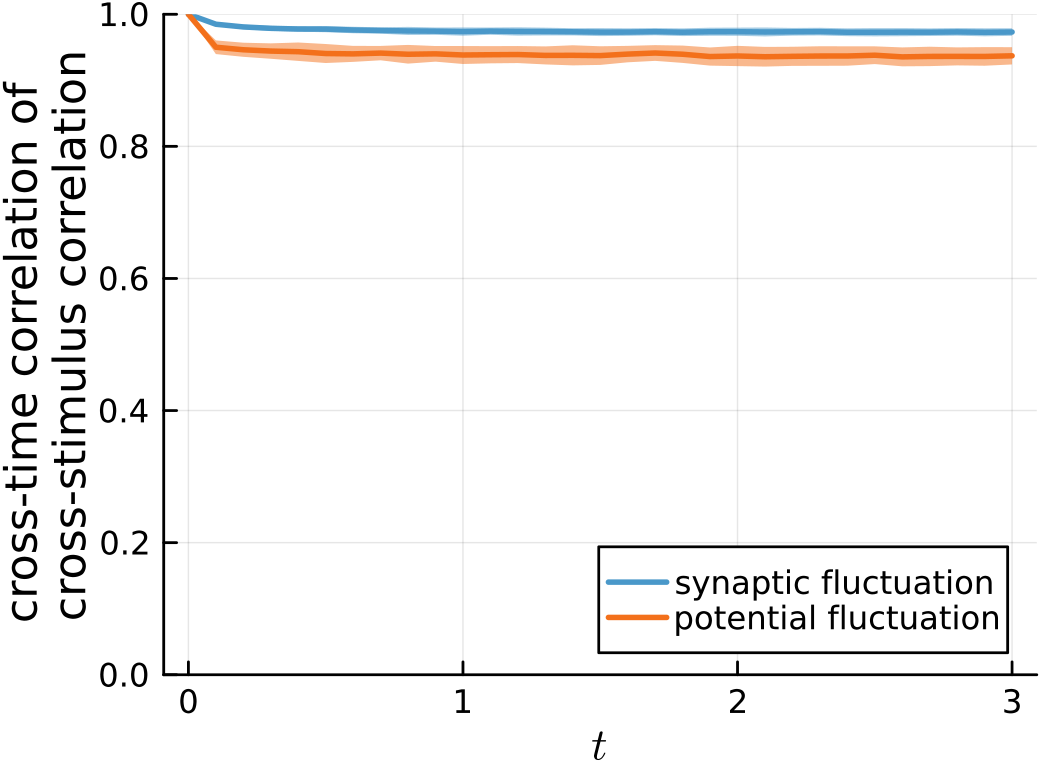
Cross-stimulus correlation matrices remain stable under synaptic and potential fluctuations. Correlations between cross-stimulus correlation matrices at *t* = 0 and *t* ∈ [0, 3*τ*] are preserved under both synaptic fluctuations (blue) and potential fluctuations (orange). Line and shaded region indicate mean and standard deviation across 100 independent simulations. The fluctuation strength is *σ* = 1.0.

**Figure S4:**
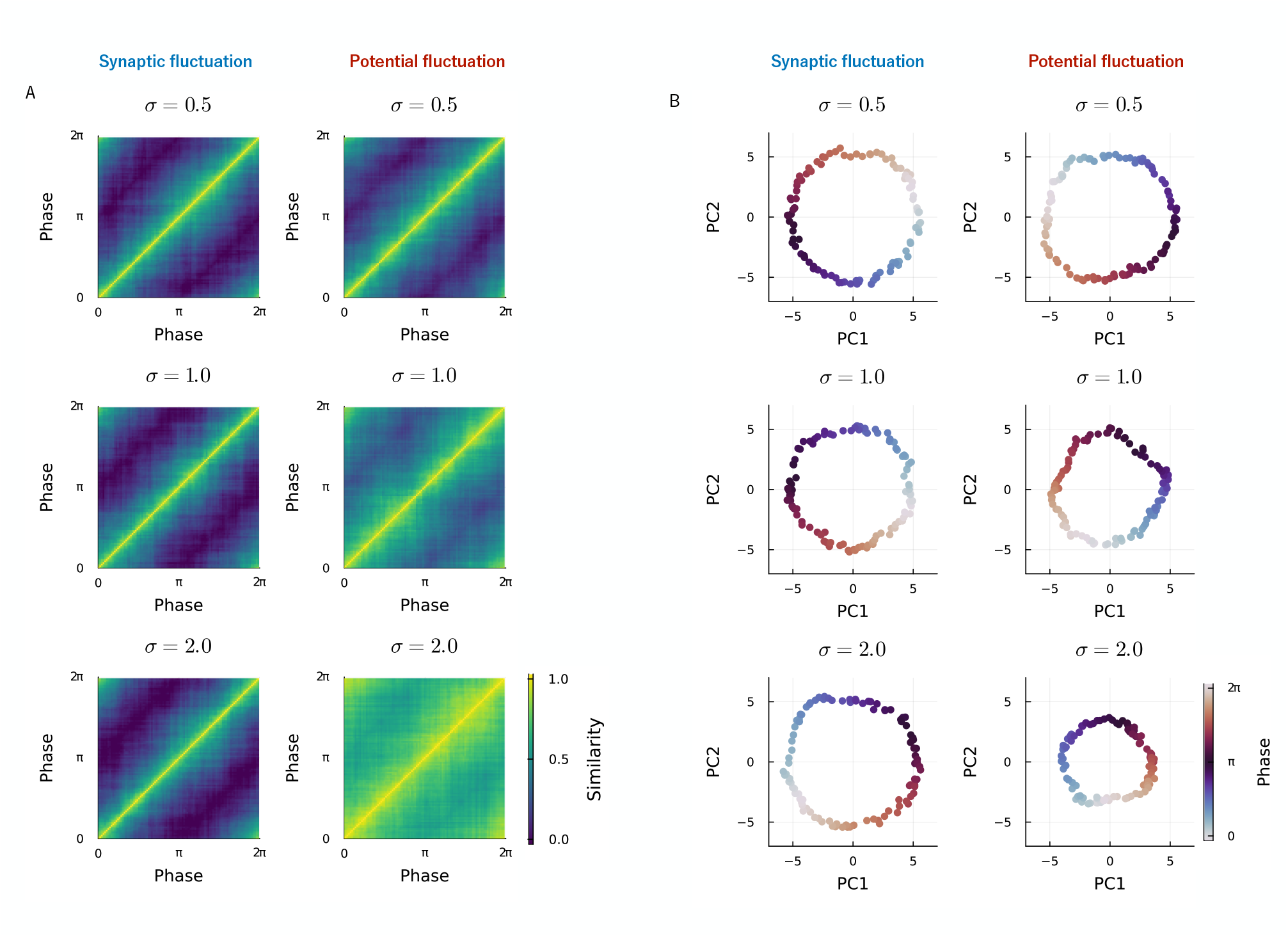
Differential effects of synaptic and potential fluctuations on representational structure are temporally consistent. (A) Cross-stimulus correlation analyses at *t* = 0 shows that potential fluctuations increase similarity across stimuli and reduce discriminability, while synaptic fluctuations preserve it. (B) PCA projections of stimulus representations at *t* = 0 for increasing fluctuation magnitudes (*σ* = 0.5, 1.0, 2.0). The ring size shrinks as *σ* increases under potential but not synaptic fluctuations. These patterns at *t* = 0 match those observed at *t* = 3*τ* (Fig. 3), confirming temporal consistency.

**Figure S5:**
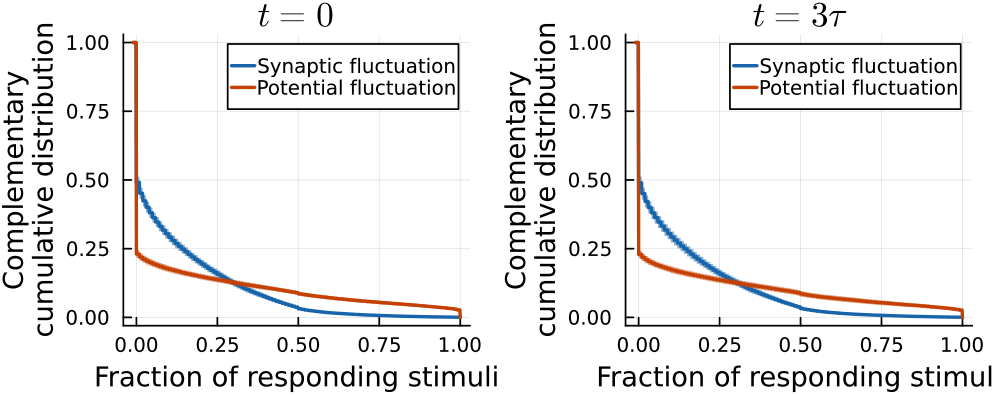
Differential effects of synaptic and potential fluctuations on fraction of responding stimuli are temporally consistent. Distribution of fraction of responding stimuli at *t* = 0 (left) and *t* = 3*τ* reveals the consequence of distinct tuning dynamics is temporally consistent: potential fluctuation produce both broadly tuned neurons and non-responsive neurons more than synaptic fluctuation. Curves and shaded regions indicate mean and standard deviation over 100 simulations. The fluctuation strength is *σ* = 2.0.

**Figure S6:**
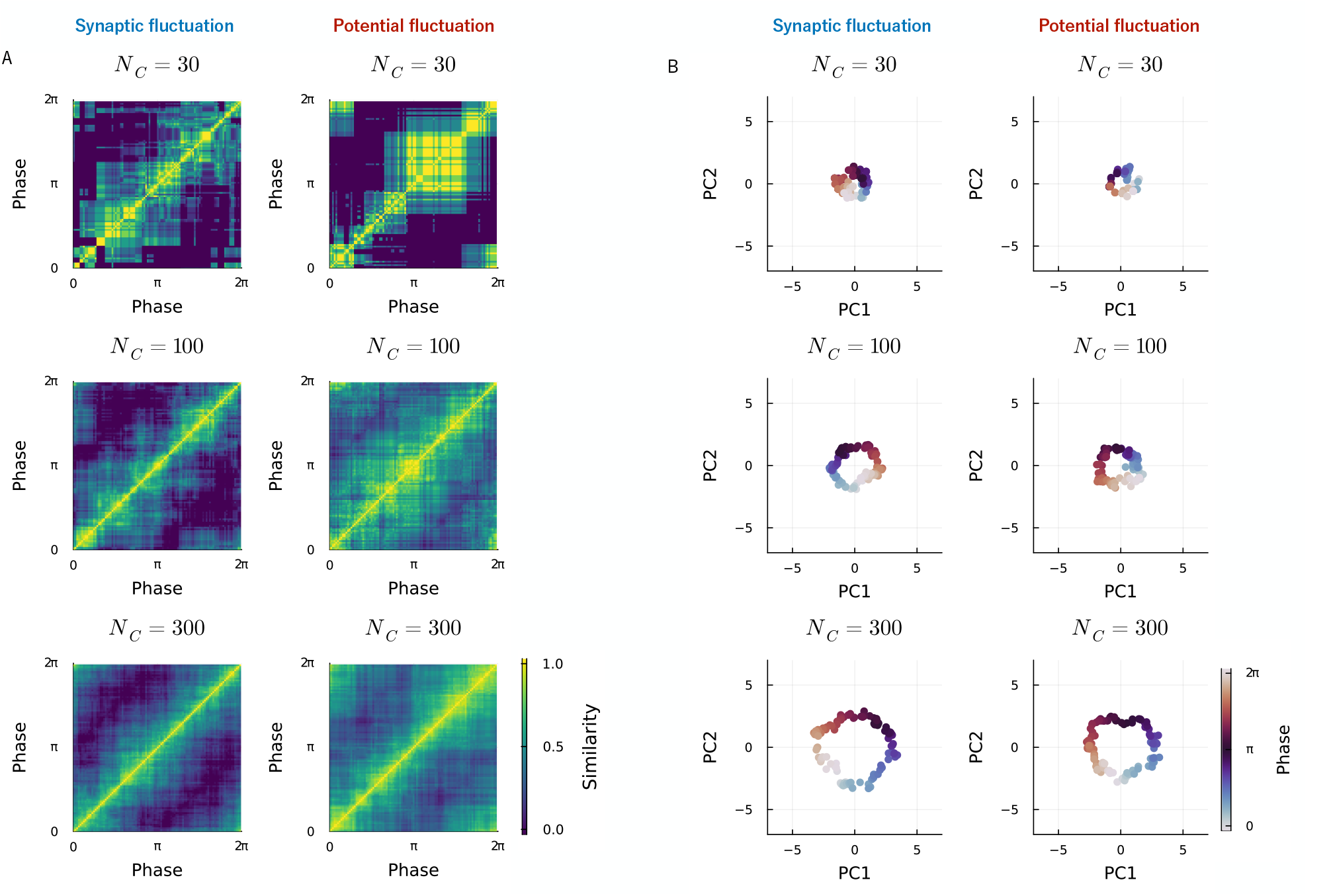
Collapse of the representational structure with small numbers of hidden neurons *N*_*C*_. (A) Cross-stimulus correlation analysis for small number of hidden neurons (*N*_*C*_ = 30, 100, 300) shows that the similarity structure collapses as *N*_*C*_ decreases under both synaptic and potential fluctuations. (B) PCA projections of stimulus representations for small number of hidden neurons (*N*_*C*_ = 30, 100, 300) show that the ring size shrinks as *N*_*C*_ decreases under both synaptic and potential fluctuations. The results shown are at *t* = 3*τ* .

**Figure S7:**
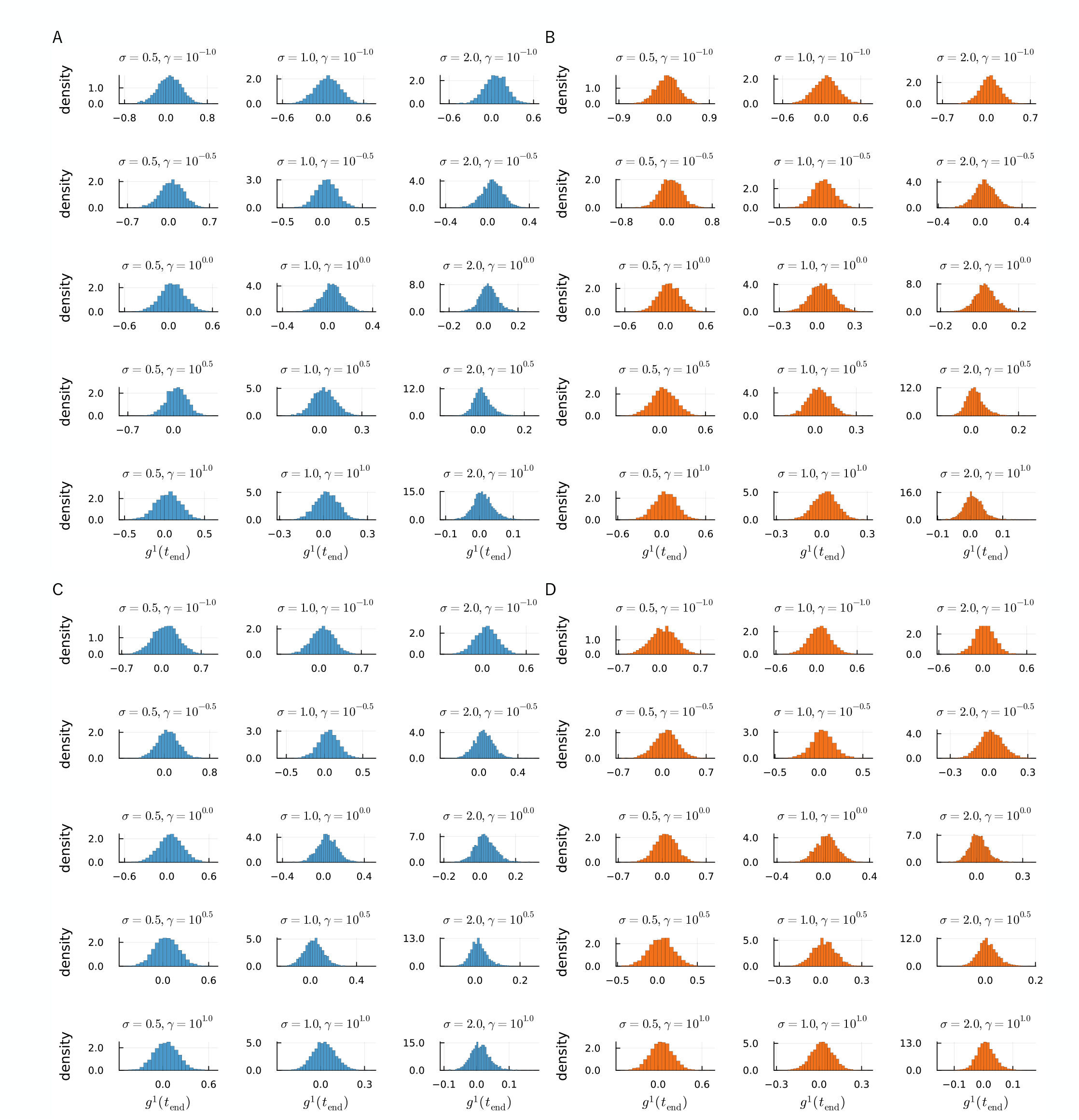
Histogram of the product of linear readout and valence *g*^*m*^(*t*) = *y*^*m*^(*t*)*L*^*m*^(*t*) at the timing of test *t* = *t*_end_ under learning of a single (A,B) or two (C,D) stimulus-valence pairs with synaptic (A,C) and potential (B,D) fluctuation. Parameters of the network were *f* = 0.1, *N*_*S*_ = 100, *N*_*C*_ = 1000 with the designated values of *σ* and *γ*.

**Figure S8:**
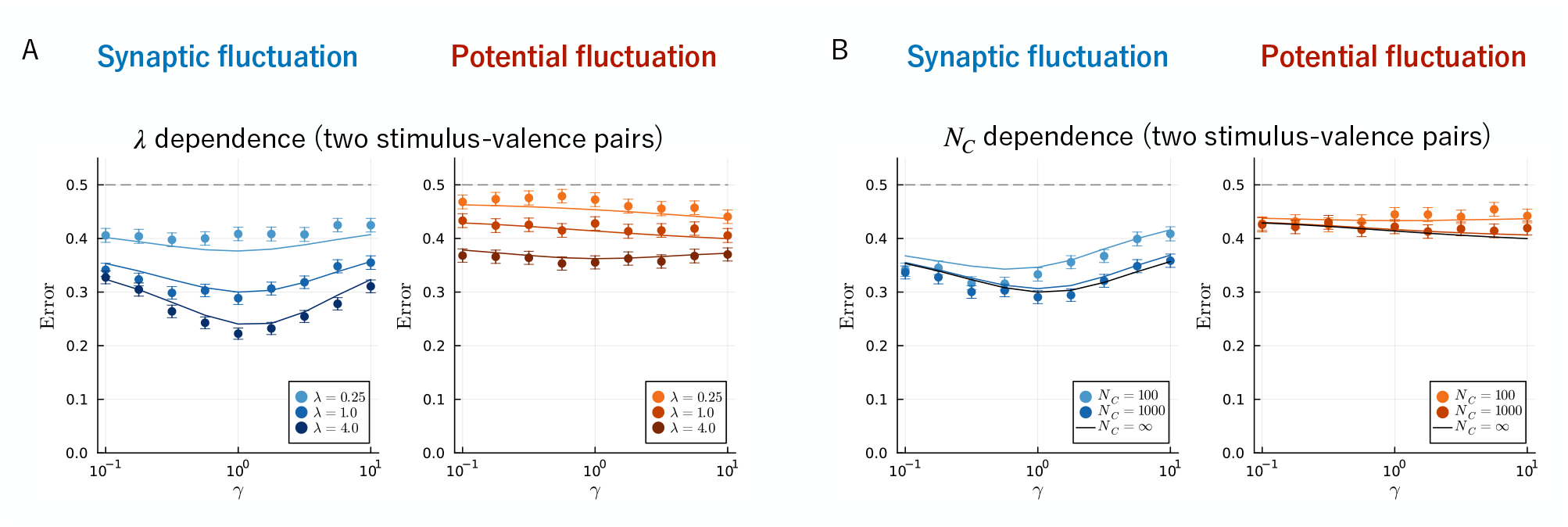
Robustness of analytical approximation and optimal values of *γ* to the increase and decrease of *λ* and *N*_*C*_, respectively. (A) *γ*− and *λ*− dependence of classification error for learning of two stimulus-valence pairs. (B) *γ*− and *N*_*C*_− dependence of classification error for learning of two stimulus-valence pairs. Points and error bars indicate the mean and the bootstraped standard deviation of 3000 independent simulations. Solid curves indicate the analytical expression (Eqs. (7) and (13) with the formula of 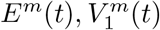, and 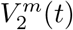 set by Eqs. (9), (11), and (12) for (A) and Eqs. (16), (17), and (18) for (B). Parameters of the network were *f* = 0.1, *N*_*S*_ = 100, *N*_*C*_ = 1000, *r* = 0.5[unit time^−1^], *σ* = 2.0, *λ* = 1.0[unit time^−1^], unless otherwise indicated in each panel.

